# A simple agent-based hybrid model to simulate the biophysics of glioblastoma multiforme cells and the concomitant evolution of the oxygen field

**DOI:** 10.1101/2023.11.27.568917

**Authors:** Luis Saucedo-Mora, Miguel Ángel Sanz, Francisco Javier Montáns, José María Benítez

## Abstract

Background and objectives: Glioblastoma multiforme (GBM) is one of the most aggressive cancers of the central nervous system. It is characterized by a high mitotic activity and an infiltrative ability of the glioma cells, neovascularization and necrosis. GBM evolution entails the continuous interplay between heterogeneous cell populations, chemotaxis, and physical cues through different scales. In this work, an agent-based hybrid model is proposed to simulate the coupling of the multiscale biological events involved in the GBM invasion, specifically the individual and collective migration of GBM cells and the concurrent evolution of the oxygen field and phenotypic plasticity. An asset of the formulation is that it is conceptually and computationally simple but allows to reproduce the complexity and the progression of the GBM micro-environment at cell and tissue scales simultaneously. Methods: The migration is reproduced as the result of the interaction between every single cell and its micro-environment. The behavior of each individual cell is formulated through genotypic variables whereas the cell micro-environment is modeled in terms of the oxygen concentration and the cell density surrounding each cell. The collective behavior is formulated at a cellular scale through a flocking model. The phenotypic plasticity of the cells is induced by the micro-environment conditions, considering five phenotypes. Results: The model has been contrasted by benchmark problems and experimental tests showing the ability to reproduce different scenarios of glioma cell migration. In all cases, the individual and collective cell migration and the coupled evolution of both the oxygen field and phenotypic plasticity have been properly simulated. This simple formulation allows to mimic the formation of relevant hallmarks of glioblastoma multiforme, such as the necrotic cores, and to reproduce experimental evidences related to the mitotic activity in pseudopalisades. Conclusions: In the collective migration, the survival of the clusters prevails at the expense of cell mitosis, regardless of the size of the groups, which delays the formation of necrotic foci and reduces the rate of oxygen consumption.

## 1 Introduction

Glioblastoma Multiforme (GBM) is one of the most aggressive and common primary tumors of the central nervous system (CNS) [3, 58]. It is mainly developed in the brain hemispheres, the cerebellum, and the brain stem by an abnormal and uncontrolled proliferation of astrocytic glial cells [61, 3, 58]. In 90% of cases, GBM is produced through tumorigenesis processes from normal astrocytes [61]. The World Health Organization classification of the CNS tumors describes glioblastoma as an IDH-wildtype diffuse and astrocytic glioma with necrosis or neovascularization, or amplification of the EGFR (epidermal growth factor receptor) gene or the combination of gain of chromosome 7 and loss of chromosome 10 (+7/-10) or TERT promoted mutation [44]. Histologically, GBM is characterized by a wide and diffusive infiltration of the glioma cells in the brain, by noticeable mitotic cell activity and by neovascularization and/or necrosis [3].

Due to the motility of the GBM cells, the total extirpation of this kind of tumor is almost impossible since cells may have been previously disseminated in the adjacent healthy brain tissue enabling the tumor recurrence [31]. Thus, resection is combined with radiotherapy and adjuvant chemo- and immunotherapy as treatment of the GBM [61, 3]. Despite this, the inherent tendency of GBM cells to invade the normal brain parenchyma limits the efficiency of this kind of treatment and makes almost impossible a cure for GBM patients [3]. Evidence of that is the fact that the 5-year survival rate is only 4.7% in GBM patients [26].

The complexity of the GBM cells invasions is related to the high complexity of the process of GBM evolution which involves different cell populations with different genes and phenotypes, extracelular matrix, chemotactic gradients and physical cues which turn on a complex and dynamic tumor micro-environment (TME) with multiple interactions [34]. The tumor cell invasion can be described into four steps, namely: (1) cells detachment, (2) cell-ECM adhesion, (3) ECM deterioration and (4) migration [3, 50]. In the first three steps, some biological actions are triggered, such as the suppression of some molecules (integrins) that enable the cell-cel and cell-ECM adhesion and the secretion of proteases to degrade ECM constituents. Cells migration is the response of the cell, moving from a point to other, to specific chemical or mechanical signals [3].

### 1.1 GBM migration mechanisms

GBM cells have a natural tendency to migrate similar to glial progenitors during the development of the CNS [50]. On the other hand, the migration of the GBM cells is also linked to the phenotypic plasticity. Cell phenotypes can have a stochastic origin or be the result of a natural adaptation of the cells to their micro-environment [51, 41]. Thus, a cell can switch its phenotype from epithelial (that represents the cell-cell adhesion in primary tumors) to mesenchymal (migratory), or vice versa, as a response to micro-environmental signals [3]. Experimental evidences have proved the low mitotic capability of the migratory cells and the quiescent behavior of the proliferative ones in GBM, melanoma or breast cancers [67, 32]. This duality is known as the “go-or-grow” dichotomy [3, 67] and results in two mutually exclusive phenotypes, namely migratory and proliferative. Which phenotype prevails will depend on the natural response of the GBM cells to their micro-environment conditions such as local cell density and/or hypoxic conditions since they can trigger or avoid the GBM cell migration [3, 8, 25, 38, 2, 59, 14].

The density of cells plays an important role in the migration of GBM cells and gliomas growth [59, 57] due to it may mark the onset of tumor invasion when the cell density reaches a critical value [24]. It may also govern the ratio of migratory and proliferative cells in gliomas since cell motility will be conditioned by how crowded the local micro-environment of the cell is [59]. Cell density is also important in the transition to collective migration in glioma cells [23]. In a wide range of tumors, and specifically in GBM, the collective migration is usually the predominant one [42, 23, 63, 40]. This collective behavior is based on the physical interactions of the cells with the TME, specifically the cell-cell and cell-ECM interplays. In these interactions, leader cells are sensitive to mechanical and chemical signals and exert a great influence in the motility of their follower cells which may not have any chemical cues or cell recognition, [42, 23, 22]. Some experimental researches with different cell phenotypes (intra- and intertumoral heterogeneity) have shown that cell invasion is produced by grouping cell populations which highlights the importance of the tumor heterogeneity in GBM invasion [3].

Another signal that can trigger the GBM cell migration is the hypoxia conditions in the TME [25]. A TME is said to be under ”modestly hypoxia” if the oxygen concentration in this area is between 0.6% and 2% [4]. An exacerbated proliferation of cancer cells and a poor conformation of a pathological new vasculature, as occurs in GBM, reduce the level of oxygen available in the TME, resulting in hypoxic conditions [9]. As the oxygen is depleted, cells migrate towards areas with more oxygen available [8] which explains in part the GBM grade of malignancy. Hypoxia also promotes the survival and proliferation of cancer stem cells of GBM by clonogenesis, duplicating the population of this type of cells [9]. In this situation, the oxygen consumption will also be drastically increased, fostering cell migration.

Despite the migratory capabilities of glioma cells, GBMs exhibit necrotic areas created, most of them, by microthrombosis [29, 9, 16]. These necrotic foci are surrounded by hypercellular regions named pseupalisades which make up one of the hallmarks of GBM according to the World Health Organization [16, 9, 54]. Pseupalisades are highly hypoxic zones related to glioma cell migration since GBM hypoxic cells actively migrate from the necrotic cores towards areas with more oxygen available [3, 17, 9, 16]. Mechanisms that cause these highly dense population of pseudopalisades are not clear [17]. Initially, the necrotic core conformation was thought to be produced by high proliferation of tumor cells in zones away from the vascular supply of oxygen [17]. However, it has been proved that pseudopalisade cells are between 5% and 50% less proliferative and 6 to 20 times more apoptotic than adjacent tumor areas, [54, 16]. That means that cell accumulation is not produced by high tumor astrocytes proliferation or by inhibition of apoptotic mechanisms. The same authors performed hystological studies in GBM and conclude that the necrotic core is produced by a vaso-occlusion and intravascular thrombosis [54, 17, 16].

These complex multiphysics processes entail multiple scales and involve a continuous interaction between cells and their micro-environment. A deeper understanding of this tumor dynamics is paramount to advance in the development and in the tailoring of new prognosis and therapeutics strategies for GBM. Addressing this complex task requires a multidisciplinary research with a continuous interplay between the mathematical modeling of the tumor processes and clinical and radiological data [64, 28, 3, 5].

### 1.2 Mathematical modeling of glioblastoma multiforme

In this section, a brief description of the mathematical modeling of the GBM will be performed, including some formulations relevant to the present proposal. Interested readers could see the reviews in references [3, 64, 28, 5, 49, 35, 45], where they will be able to find examples of other formulations.

#### 1.2.1 Continuum formulations

Continuum formulations have brought important insights into GBM dynamics. The GBM biophysics is modeled through reaction-diffusion equations consisting in partial differential equations (PDEs) formulated in terms of variables such as concentration of oxygen, cell density, angiogenic proteins and enzymes [64].

In the first models, Tracqui et al. [60] formulated a system of PDEs in terms of the cell density of two cell populations during the infiltration and growth of GBM. Kim et al. [39] included the cell-cell adhesion explaining the pattern of collective migration in GBM cells and Pham et al. [52] studied the influence of the cell density in the phenotypic switch from proliferative to migratory, and vice versa, through a mathematical model based on the”go-or-grow” dichotomy. Martínez et al. [47] included the interplay between vasculature, oxygen distribution and hypoxic, normoxic and necrotic phenotypes in the dynamics of the pseudopalisades. Alonso et al. [2] adopted the”go-or-grow” dichotomy to simulate the coupling between the GBM cell invasion and the vasculature evolution. The authors modeled the oxygen concentration proportionally to the functional vasculature and considered hypoxic and normoxic phenotypes. More recently Conte and Surulesco [21] studied GBM invasion considering implicitly the oxygen supply through the evolution of the endothelial cells since they are the responsible for the nutrients supply and therefore of the phenotypic switch.

This continuum approach is suitable to study cancer biophysics at tumor scales. However, this framework is not sensitive to other important biological process underlying this scale such as cell-cell and cell-ECM interactions or vascular sprouting during angiogenesis [64]. At these cell scales, discrete models represent an alternative to the continuum formulation to study tumor dynamics, in which, furthermore, these interactions may be implemented in a considerably simpler manner.

#### 1.2.2 Discrete and hybrid models

Cellular automaton (CA) and agent-based models (ABMs) are the two principal discrete models used to mimic biophysics processes in GBM [64]. In CA, a group of cells is modeled through the same rules which are based on the conditions of their surroundings cells so that they evolve synchronously along a certain period of time. In ABMs each single cell is considered as an independent agent that evolves autonomously depending on its interactions with the rest of cells and on its micro-environment [64]. Although some CA models have been formulated to reproduce cell plasticity on tumor initiation and persistence [15] and proliferation and/or migration in GBM cells [14, 59, 67, 7], this kind of formulations may not properly reproduce the heterogeneity and the multiple interaction between cells and their micro-environment given in the GBM. For that, ABMs constitute an interesting framework to model the complex biophysics of the GBM [64] such as the growth dynamics in spheroids [27], the differences found in the velocity of normoxic and hypoxic glioma cells in collective migration [38], the influence of cell-cell adhesion in the propagation front of glioma cells [37] or the influence of the ECM on vasculogenesis [20].

The main restriction to the discrete modeling is that its application is limited to a relatively few population of cells, when compared to actual tumors, due to the computational cost [64]. However, advances in parallel processing, high performances GPU-based codes, the new generation of high performance supercomputers and the sinergy with continuum formulations are increasingly allowing to reproduce multiphysics processes of tumors [5].

Multiscale hybrid models integrate the continuum and discrete formulations bringing the best of both approaches. This type of models enables the simulation of tumor dynamics at subcellular, cell and tumor levels, with a continuous connection and interplay between all the scales [64, 28].

This framework has enabled the modeling of the phenotypic switch in brain tumors through a genprotein interaction at subcellular level [6, 66]. In [66], the authors integrated in their model the cell cycle module developed by Alarcón et al. [1] including the hypoxia as external condition. Caiazzo and Ramis [19] formulated a multiscale model to analyze the pseudopalisades formation in GBM. They modeled the biophysics characteristics of the cells through an individual force-based model whereas with the continuum part accounted for the diffusion of oxygen in the tissue. Recently, Sadhukhan et al. [56] developed a forced-based hybrid model to study avascular tumor growth through protein/gen, cell and tissue scales. The diffusion of oxygen and nutrients was formulated by Fick’s law. Sadhukhan and Mishra [55] enlarged this model to study cell heterogeneity produced by phenotypic changes during avascular tumor growth. In both studies, the authors considered five possible phenotypes at cell scale: hypoxic, proliferative, apoptotic, necrotic and quiescent. Recently, Benítez et al. [12] developed a three-dimensional hybrid ABM to simulate the coupling between tumor induced angiogenesis evolution and oxygenation of GBM cells. The diffusion of both the oxygen and the VEGF was formulated by means of Fick’s law. Cells and vasculature were modeled as individual agents.

Despite the efforts made in the last twenty years, some of the processes involved in the development of GBM are not fully understood. This is evidenced by the low rates of survival in the first five years since diagnosis (less than 5%) and the fact that the median survival of GBM patients is 6 to 16 months [26, 58], so some advances in biophysical modeling are needed. Such is the case of modeling and understanding the collective migration of cells [30, 23, 42], the biochemical and biophysics processes in the TME [56], the need to account for other phenotypes than the proliferative and migratory phenotypes in GBM or to analyze the consequences for invasion [3]. Furthermore, not many of the researches in this field have addressed the simulation of the coupling between the evolution conditions at tissue level and the evolution of the cell events so that more insight in this issue is needed [3].

The present work intends to shed light on these issues by means of a simple 3D agent-based hybrid model capable of simulating the coupled process of individual and/or collective cell migration, the concomitant evolution of the oxygen field in the ECM and phenotypic plasticity, considering five phenotypes: migratory, quiescent, proliferative, hypoxic and necrotic. The simulation through different scales, macroscopic for the oxygen evolution and microscopic for cell events, enables the visualization and analysis of the mutual influence of both scales during GBM cell migration. This approach also allows to reproduce some other hallmarks in GBM such as the formation of the necrotic cores and pseudopalisades. The simplicity of the model is due to the use of few genotypic and biophysical parameters. This simplicity and easiness for parallelization turns out in reasonable simulation times. Although the authors have developed a 3D multiscale angiogenesis model for GBM [12], in the present model, in order to focus on the cell response, the influence of the vasculogenesis is simulated through the evolution of the oxygen field supplied by the blood vessels at the tissue level, as it is performed in experimental tests [8].

The model has been formulated for three dimensions, however it has been applied to 2D cases in order to depict clearly the bio-physical processes in GBM and due to the fact that many in vitro tests are performed in 2D petri dishes or 2D microfluidic devices. Although the model has been developed to reproduce GBM hallmarks, these biological processes can be also simulated for other types of tumors. The model has been applied to benchmark cases and to reproduce in vitro experimental tests.

The paper is structured as follows. In Section 2, the model is described from a biophysical viewpoint. In Sections 3 and 4, the mathematical formulation of the model and an analysis of the parameters sensitivity are respectively addressed. Some applications, including the simulations of experimental tests, are shown in 5. In Section 6, the model limitations are decribed and the future work is outlined. Finally, some conclusions are stated in Section 7.

## 2 Genotypic and phenotypic modeling of cell migration

Although the present model is based on the “go-or-grow” dichotomy, we have reformulated it to the “go-or-stay” one in order to consider cell phenotypes other than the proliferative or migratory ones. We replace *grow* by *stay* because we assume that proliferation (growth) is produced after a period of quiescence and when suitable conditions of oxygen and cell density are given. If these conditions are not produced, a quiescent cell might migrate or go into hypoxia and even die, resulting in five possible phenotypes as the result of the interaction between the cell and its micro-environment, namely: quiescent, migratory, hypoxic, proliferative, and necrotic.

In this proposal, the influence of the micro-environment in the migration of a given cell is characterized by the cells that surround it and by the oxygen available in the extra cellular matrix. Although the model has been focused on the chemotaxis induced by oxygen and on the influence of the cell density, its structured design easily allows for the introduction of new substances, including drugs, other nutrients such as glucose and the mechanical forces between cells, which have not been considered so far in this study.

The migration of the cells is simulated along a certain period of time which is divided into time steps, ∆*t*. In every time step, the evolution of the oxygen field and the cell density distribution are computed in two different layers continuously interplaying. In a first layer, the oxygen concentration in the tissue is calculated solving the diffusion equation by a finite element code. In the second layer, an ABM is implemented as a game of perfect information since it is based on the game theory. It consists of a tumor cell population distributed within the elements of finite element mesh automatically generated by the finite element algorithm. Each element of the mesh makes up the micro-environment of a cell *i* placed inside of it. This micro-environment, or element, is denoted by *α*_*i*_. At each time step, the cells are defined as individual agents characterized by their position in the mesh, through their position vectors, their diameter and their phenotypic and genotypic parameters. Genotypic variables can be obtained from experimental tests, whereas the phenotypic ones are estimated.

### 2.1 Genotypic parameters

Two types of genotypic parameters are considered. The first group concerns the individual behavior of the cells from the oxygen concentration viewpoint. The second one accounts for the cell behavior with respect to the presence of other cells in its micro-environment and their tendency to form clusters.

The behavior of a single cell *i*, at time *t*, in an oxygen field is modeled by the maximum time that it can be in anoxia, *t*_*a*_, the minimum level of oxygen needed to survive in normoxia, 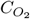, its average oxygen consumption rate, 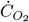, and the minimum time needed to produce cell proliferation, *t*_*m*_. Although this model does not address growth process itself, tumor growth has been simulated through a mitosis process consisting in a cell duplication after a time, *t*_*m*_, in a normoxic micro-environment and a previous quiescent phenotype. The structure of the model easily allows a more sophisticated implementation in the future. Other variables have been considered to model how the cell perceives the oxygen field. These genotypic parameters are *A*_1_ and *B*_1_ intervening in Equation (4) below. As it will be explained in detail in Sections 3.2.1 and 4, *A*_1_ and *B*_1_ model the cell behavior for low and high oxygen concentration.

The influence of surrounding cells on an individual cell and on an other group of cells is reproduced through the optimum biological cell density in a cell micro-environment, *ρ*_*op*_, and a parameter of null cell density, 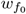. Cell density is defined as the ratio of the volumes of all individual cells within an element of the mesh and the volume of that element. Hence, it should be interpreted as a relative density or mass ratio, without units. If this density equals one, the element is fully occupied by cells. *ρ*_*op*_ accounts for the maximum density of cells in an element whereas 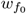 takes into consideration how comfortable the cell is under isolation. By means of these density variables, collective behavior of cells can be simulated as it will be explained in Section 3.2.2.

Another genotypic parameter included in this model is the average velocity of each cell during its displacement, *v*_*i*_. In Table 1, a summary of the genotypic parameters is shown.

**Table 1:** Summary of the genotypic parameters.

| Parameter | Description |
| --- | --- |
| $t_a$ | Maximum time that a cell $i$ can be in apnea |
| $C_{O_2}$ | Minimum oxygen concentration needed to survive (switch from normoxia to anoxia) |
| $\dot{C}_{O_2}$ | Oxygen consumption rate of the cell |
| $t_m$ | Minimum time needed to produce cell proliferation |
| $A_1$ | Cell behavior for low oxygen concentration |
| $B_1$ | Cell behavior for high oxygen concentration |
| $\rho_{op}$ | Optimum biological cell density |
| $w_{f_0}$ | quiescent trend of the cell when it is by its own |
| $v_i$ | Average velocity of the cell $i$ during displacement |

### 2.2 Actions (decisions) and phenotypic plasticity

Once the interactions between the cells and their micro-environments have been assessed, each single cell will only take one of the following decisions, also called actions; specifically (see Table 2), a cell will migrate, Λ_*m*_, stay, Λ_*s*_, grow, Λ_*g*_, be in apnea, Λ_*a*_, or die, Λ_*d*_. The two first, to migrate or to stay, are called here primary actions since the rest of actions are derived from them. There is a particular case in which a cell can perform the actions of migrating and being in apnea simultaneously, as it will be described in Section 3.2.6.

**Table 2:** Summary of the actions and their relation with phenotypes.

| Action | Description | Phenotype | Description |
| --- | --- | --- | --- |
| $\Lambda_m$ | Migration | $f_m$ | Migratory |
| $\Lambda_s$ | Stay | $f_q$ | Quiescent |
| $\Lambda_g$ | Grow | $f_p$ | Proliferative |
| $\Lambda_a$ | Be in apnea | $f_h$ | Hypoxic |
| $\Lambda_d$ | Die | $f_n$ | Necrotic |

Actions and cell phenotypes are related. From a formulation viewpoint, we assume that, initially, every cell has a natural tendency to *go* or to *stay* given by a stochastic intrinsic phenotype, namely migratory or quiescent. However, this intrinsic phenotype may change because the cell micro-environment forces the cell to perform a different action from migrating or staying. For example, a cell with a migratory intrinsic phenotype may be forced to execute the action of *staying* because there is no space to move in its surrounding due to a high cell density. Thus, the cell phenotype will change from migratory to quiescent because of the environment. In this case, its survival will depend on the oxygen available in the zone. If there is not enough oxygen, the cell phenotype will switch to a hypoxic one where the cell is in apnea. Even more, if a certain cell is under hypoxic conditions for a period of time longer than its genotype allows, its phenotype will change to necrotic. In the same way, a quiescent phenotype can switch to migratory one if there is not enough oxygen and/or there is a great accumulation of cells in the cell location. Finally, quiescent cells with enough oxygen and suitable cell density in their surroundings will change its phenotype to proliferative after a period of time in quiescence. Therefore, we considered five phenotypes associated to the above mentioned actions, namely, migratory *f*_*m*_, quiescent, *f*_*q*_, proliferative *f*_*p*_, hypoxic *f*_*h*_ and necrotic *f*_*n*_. In Table 2, the relationships between the actions and the phenotypic that a single cell can adopt in a certain time step are summarized.

## 3 Mathematical formulation

As above mentioned, the model is composed by two layers, continuum and discrete, which respectively contain the information of oxygen concentration and cells density of the cell micro-environment, with the key features that lead the cells towards their decisions. As a consequence of these actions, the tumor micro-environment will change. For example, when a cell migrates, the density of cells at the provenance zone will decrease whereas the one at destination will increase. Simultaneously, the consumption of oxygen will be increased in the region of destination due to the extra “sink”. This requires a continuous interplay between the continuum and discrete layers of the model.

Two codes have been developed in Julia programming language for both the continuum and the discrete layers. In the former, the oxygen diffusion equation is solved in every time step by the Finite Element Method (FEM), and in the latter, an ABM is executed at the same time step. In the ABM, a certain number of cells are introduced with an initial phenotype as will be explained in Section 3.2. The cell decision is individual without accounting for the rest of cells, so there is no specific order in taking the decision which makes the ABM code be flexible. The cell decisions in one time step will be executed in the following one. Thus, in every time step, the diffusion equation is solved with the new positions of both the migratory and the new cells, resulting from the decision process of the previous step. These new positions are considered new sinks for the FEM code. Once the diffusion problem is solved, cells make their decisions individually and according to the new oxygen field and the current cell arrangement. This decision imply a new cell arrangement that will be executed in the next time step.

### 3.1. Continuum formulation

The oxygen concentration in each point of the region is provided by a FE model that solves the oxygen diffusion equation. In this formulation, angiogenesis is not directly considered, so no source terms are included. Instead, the effect of the vasculogenesis is mimicked by the evolution of the oxygen concentration initially supplied by the neovasculature, in a similar way as it is performed in experimental microfluidic models of GBM [8], and as will be explained in Section 5. This model is very simple and implements Equation (1).

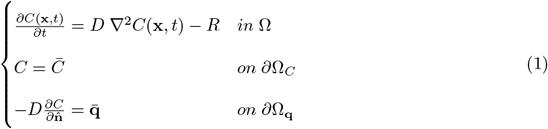

In this equation, *C*(**x**, *t*) is the oxygen concentration, **x** is the position vector of a certain point of the continuum, *t* is the time variable, *D* the diffusivity of an isotropic medium, ∇^2^ is the Laplace operator, with the meaning of the divergence of the gradient of *C*(**x**, *t*), and R is the sink term. In the boundary terms, the letter with a bar means that such magnitude is known, for example, the flux 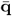 through a surface of the flux boundary *∂*Ω_**q**_ with normal unit 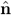. With respect to the boundary conditions adopted in this work, 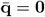 on *∂*Ω_**q**_ in most of the cases. Since oxygen diffusion is much faster than the changes in cells, it suffices to compute the stationary solution because of the difference in the time scales between continuum and discrete events, so *∂C/∂t* = 0 in Equation (1).

To solve numerically Equation (1), the first equation is multiplied by a virtual concentration and integrated over the element domain, *α*_*i*_, to obtain the Galerkin formulation of the problem. Integrating by parts, see [10], and assuming that 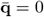, the equivalent weak formulation of the problem is obtained at element level, Equation (2).

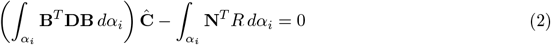

where **N** is the matrix of the element shape functions, **B** = ∇**N, D** = *D***I** is the diffusivity coefficient matrix for the isotropic case, **I** is the second order identity tensor and **Ĉ** is the nodal vector of the oxygen concentrations. The multiscale coupling is performed in the sink term which is formulated through Equation (3) assuming that the oxygen consumption, 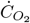, is the same for all the cells (except the necrotic ones that is null). The upscaling is carried out accounting for the number of glioma cells, except the necrotic ones, in the element of the FEM mesh, 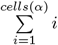.

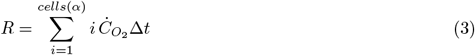

The downscaling is performed through the information of the oxygen concentration available for the cell. For that, the oxygen level is averaged in each element of the FE mesh so that all the cells inside the same element have the same oxygen concentration.

### 3.2 Agent-based formulation

In the discrete layer, the information of the cell density is given and each cell decision is performed individually. In each time step, we consider that a single cell has an intrinsic phenotype defined by its natural tendency to go or to stay regardless of the micro-environment restrictions to these actions. Determining experimentally if the intrinsic phenotype of a cell is migratory or quiescent is a difficult task, moreover considering the large number of cells involved in tumor processes. For those reasons, this natural trend is assumed to be stochastic so that it can be modeled by a normal distribution function denoted by 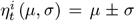. In this function, *μ* is the media, *σ* is the standard deviation and 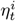 is the value of the normal distribution function given to a single cell *i* at time *t*. A low value of standard deviation is adopted to avoid high fluctuations in the phenotype. Therefore, the values of 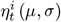 with a mean of *μ* = 0, i.e. 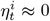, will give to most of cells a marked tendency to stay whereas those values with a media of *μ* = 1, i.e. 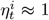, will model a clear trend to migrate. According to this, cell decision is based on determining if the value of 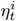 is low or high. Cell decision will not only depend on the cell intrinsic phenotype since its micro-environment conditions, characterized by the oxygen and cell density, will allow or restrain this natural tendency to stay or to go.

#### 3.2.1 Influence of the oxygen

The influence of the oxygen field on the cell behavior has to be formulated so that the model promotes the cell migration as the oxygen concentration within its micro-environment is reduced. This behavior is formulated by means of the Weibull distribution function 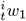, Equation (4), which represents the probability that a cell stays in its micro-environment *α*_*i*_ under a level of oxygen 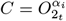. The response of the cell against the oxygen field is modulated by two parameters, *A*_1_ and *B*_1_, which respectively determine the spreading out and the shape of the function, Figures 1 and 2.

**Figure 1:**
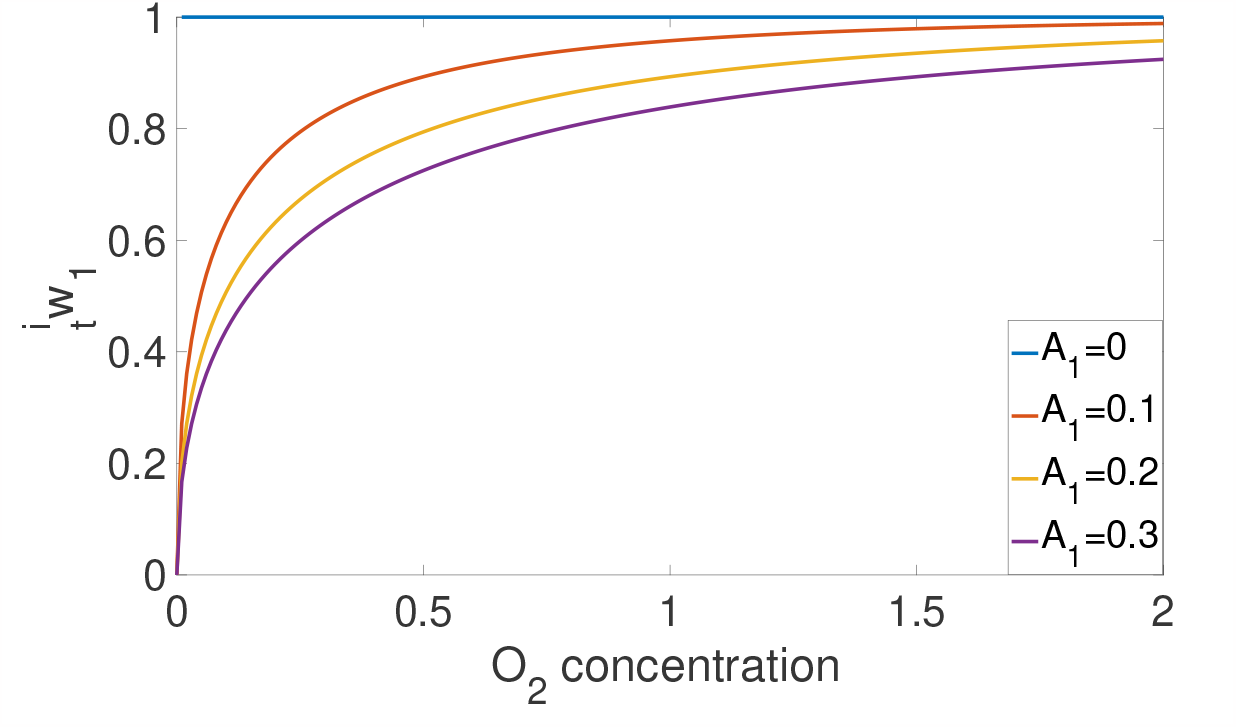
Graphical representation of 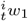 in terms of different values of *A*_1_ with *B*_1_ = 0.5

**Figure 2:**
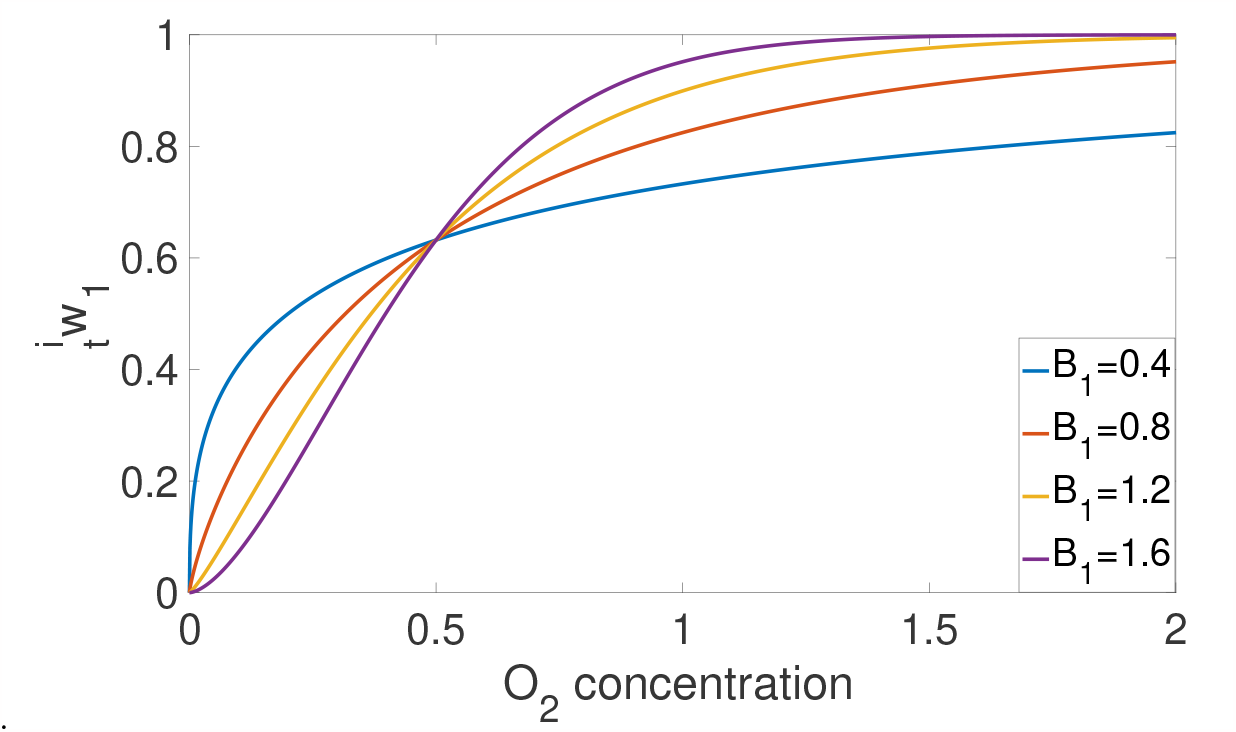
Graphical representation of 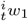 in terms of different values of *B*_1_ with *A*_1_ = 0.5

In Figure 1, the results of Equation (4) are depicted keeping *B*_1_ = 0.5 and considering four different values of *A*_1_, specifically 0, 0.1, 0.2 and 0.3. When *A*_1_ = 0, Equation (4) turns out a value of 1 for any oxygen concentration and any *B*_1_ *>* 0 which is reflected in the horizontal line in Figure 1. This means that the oxygen field in this micro-environment does not have any influence on the migratory behavior of the cell since the cell will always have the same sensitivity to low and high oxygen concentrations. In this case, the cell will have a null tendency to migrate under any level of oxygen. This behavior is lost if *A*_1_ */*= 0. The results of Figure 1 show that, for the same oxygen concentration, cells with larger values of *A*_1_ will migrate earlier than other cells with lower values of *A*_1_, Figure 1.

Consider now that *A*_1_ = 0.5 is kept constant and different values of *B*_1_ are adopted, namely 0.04, 0.08, 0.12 and 0.16. With these values, the results of evaluating Equation (4) are depicted in Figure 2. The figure shows that all the curves intersect at a point. This point corresponds in the abscissa axis to the value of *A*_1_ considered, *A*_1_ = 0.5 in this case, whereas in ordinate axis to the value of 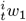 when *B*_1_ = 0, i.e. 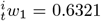. Up to this point, the probability that the micro-environment allows the cells to stay will be greater, for the same oxygen concentration, as the values of *B*_1_ is diminished. However, from this point, this tendency switches and the probability of staying is greater for higher values of *B*_1_ and for the same oxygen concentration.

Therefore, *A*_1_ and *B*_1_ modulate the response of a cell under a certain concentration of oxygen. According to this, we can state that Equation (4) properly reproduces that the probability that a cell migrates is higher as the oxygen concentration is reduced and that modeling the cell behavior, from a migratory viewpoint, in an oxygen field is performed by both *A*_1_ and *B*_1_.

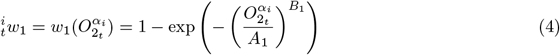

#### 3.2.2 Influence of the cell density. Individual and collective migration

The individual cellular migration triggered by the cell density *ρ*(*α*_*i*_) in a given micro-environment *α*_*i*_ is modeled following a similar formulation as in the case of the oxygen. In this case, the micro-environment promotes cell migration as *ρ*(*α*_*i*_) is high. Whether this cell density is high or not is given by the optimum density, *ρ*_*op*_. Thus, the probability that a cell migrates has to increase as the cell density is greater than the optimum density, *ρ*(*α*_*i*_) *> ρ*_*op*_. Therefore, this probability can be calculated through Equation (5). The variable adopted is the void ratio 1 − *ρ*(*α*_*i*_) instead of *ρ*(*α*_*i*_) since we look for the probability that a cell stays is equal to 1 when the cell density is null (void ratio equals to one). To properly model this behavior, the role of the shape and scale parameters is also paramount. Thus, the shape parameter has been estimated as *B*_2_ = 2 to obtain a smooth shape of the function and to force that the probability of staying is null when the cell density is close to one regardless the optimum density value. The scale parameter is 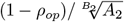. The value of the parameter *A*_2_ has been adopted as *A*_2_ = 6.9 so that, for all 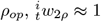 when *ρ* ≤ *ρ*_*op*_ and 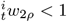 when *ρ > ρ*_*op*_.

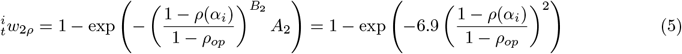

In Figure 3, the results of Equation (5) have been depicted in terms of the values of the cell densities, *ρ* = *ρ*(*α*_*i*_), in the environment of a given cell *i* and considering different options of cells optimum density, from *ρ*_*op*_ = 0 up to *ρ*_*op*_ = 1. According to this figure, the probability that the cell density in a element of the FE mesh allows a cell to stay is maximum and equal to 1 when *ρ < ρ*_*op*_. In the case of *ρ*_*op*_ = 1, this probability is always equal to 1 regardless of the value of *ρ*. For the rest of cases, i.e. *ρ*_*op*_ *≠* 1, if the cell density is greater than *ρ*_*op*_, the probability that the micro-environment allows the cell to stay decreases as the cell density *ρ* is increased. In this cases, for the same cell density, the probability diminishes when *ρ*_*op*_ is reduced.

**Figure 3:**
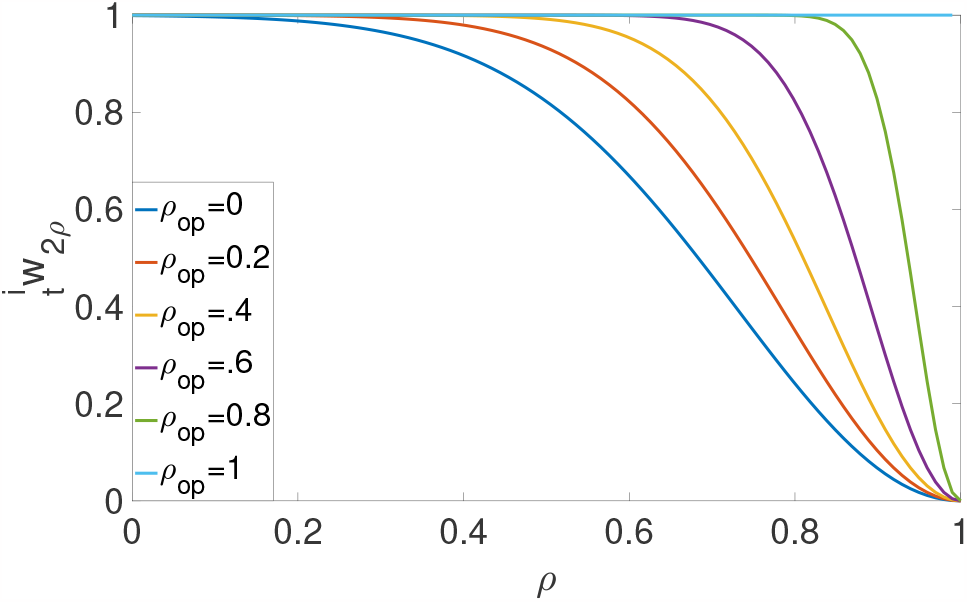
Graphical representation of 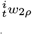in terms of different values of *ρ*_*op*_

According to the above mentioned, we can conclude that 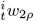 can be interpreted as the probability that a given cell stays in *α*_*i*_ in function of *ρ*(*α*_*i*_), so that if the cell density is high compared to *ρ*_*op*_, the cell will migrate individually towards areas with a lower cell density.

The collective behavior is formulated at cell scale through two parameters which avoids formulation at subcellular scale. Specifically, we assume that cell-cell interaction can be simplified considering that cells exhibit flocking: a harmonic and collective arrangement found during birds migration [11], in human stem-cells [43] and in tumors invasion [23, 30, 36]. In this formulation flocking is modeled by the tendency of cells to form clusters with high cell density. The modeling of the collective behavior is based on the formation of groups of high cell density so that the migration of the cell in this case is conditioned to reach *ρ*_*op*_ in the corresponding micro-environment. For that, Equation (5) is corrected according to Equation (6).

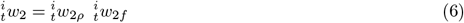

In Equation (6), 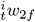 is the corrector function, also called flocking function, which has been formulated so that a cell tends to migrate as long as the cell density in the micro-environment is not close to the optimum density. For that, Equation (7) is proposed, where *w*_*f*0_ represents the tendency of a single cell to stay if it is alone, i.e. when *ρ*(*α*_*i*_) ≈ 0. The values of *w*_*f*0_ are given in the interval 0 *< w*_*f*0_ ≤ 1.

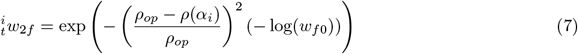

In order to understand how the correction is performed, the results of Equation (5) are depicted in Figure 4 for three values of *ρ*_*op*_ (0.4, 0.6 and 0.8). The same three values of *ρ*_*op*_ have been used to show the results of Equation (7) in Figure 5 considering *w*_*f*0_ = 0.1. Finally, the corrected function, Equation (6), is depicted in Figure 6. According to the results shown in these figures, when the optimum density has not been reached, the individual tendency of the cell to stay 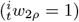 is reduced by 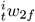whenever *ρ < ρ*_*op*_. When optimum density is reached (*ρ* = *ρ*_*op*_) then 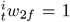 and therefore 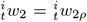. When *ρ > ρ*_*op*_, 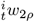 is similarly corrected, having the singularity that 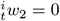 if *ρ* = 1.

**Figure 4:**
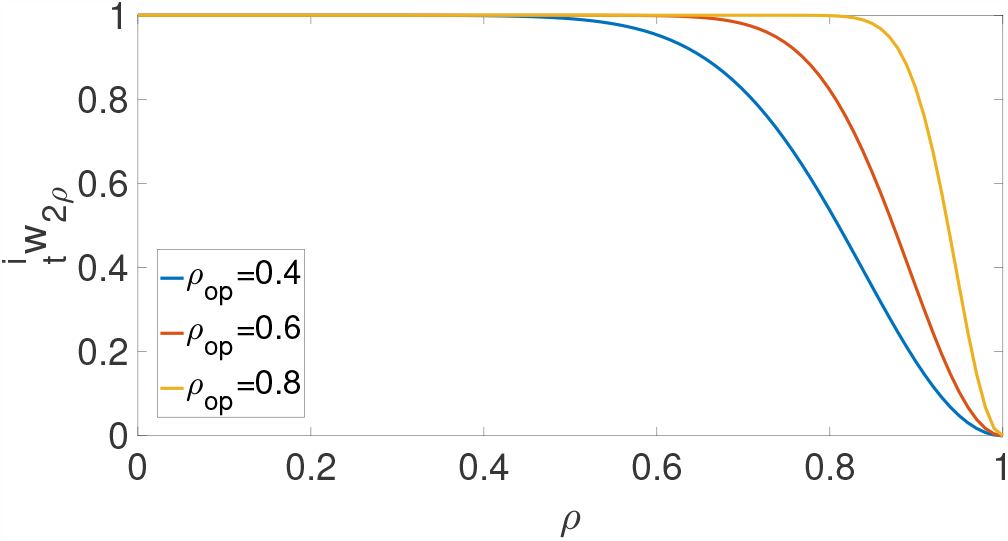
Graphical representation of 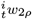 for *ρ*_*op*_=0.4, 0.6 and 0.8.

**Figure 5:**
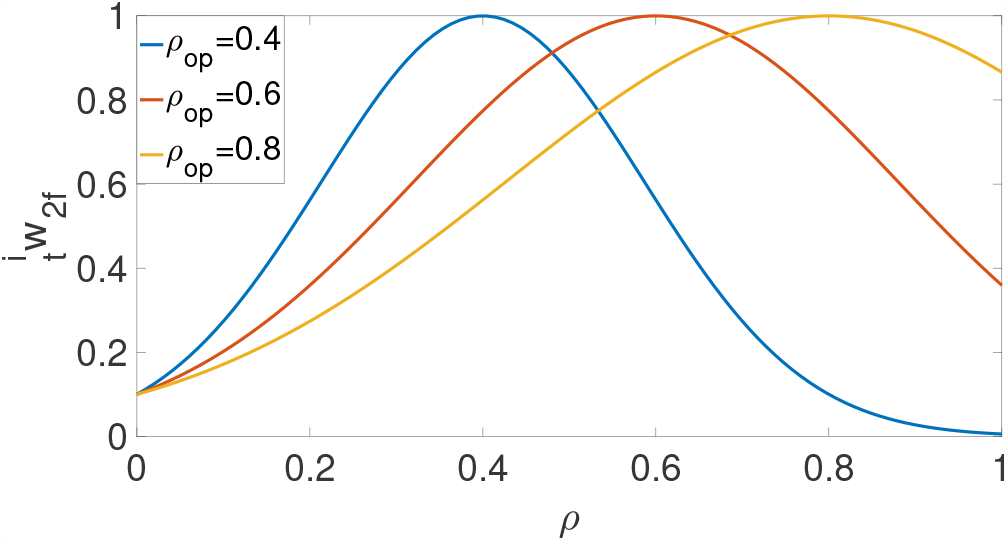
Graphical representation of 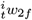 for *w*_*f*0_ = 0.1 and *ρ*_*op*_=0.4, 0.6 and 0.8.

**Figure 6:**
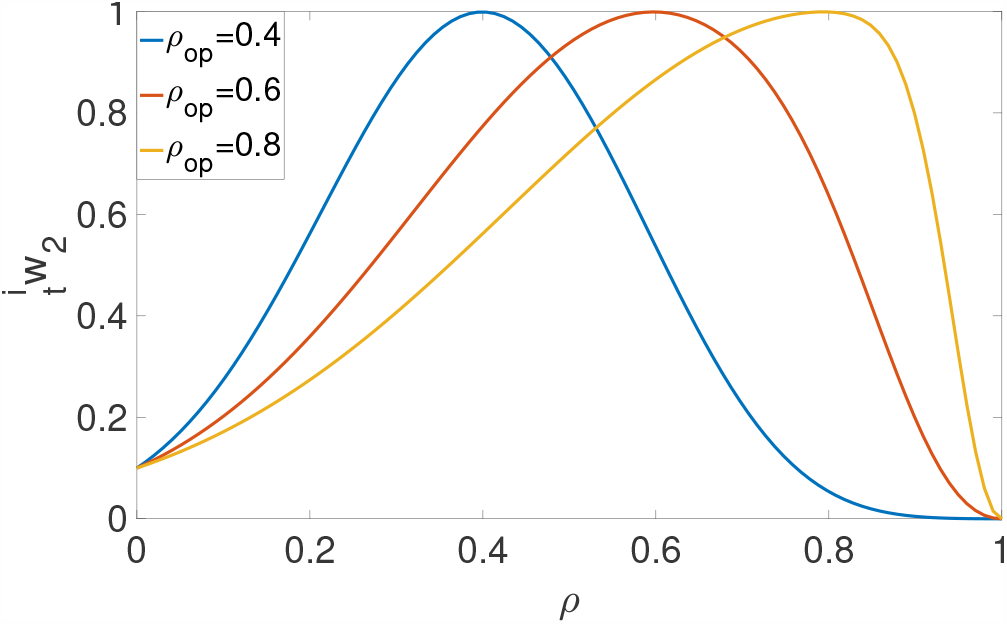
Graphical representation of 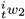for *w*_*f*0_ = 0.1 and *ρ*_*op*_=0.4, 0.6 and 0.8.

Since the Equation (6) is a corrected probability, it will only be a distribution function when there exits an individual migration, i.e. 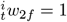 for all *ρ*(*α*_*i*_), and therefore 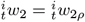 for any *ρ*(*α*_*i*_). Despite this, 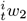 can be interpreted as the tendency of a cell staying in function of the cell density. Therefore when the cell density is zero, the cell’s tendency to stay will be modeled by the parameter *w*_*f*0_. For the rest of densities this tendency will be increased as *ρ* → *ρ*_*op*_.

#### 3.2.3 Cells migratory behavior based on the level of oxygen and the cell density

That the cell micro-environment allows a cell to keep its phenotype or forces it to change phenotype can be produced by either the oxygen field and the cell density. Thus, each cell must develop an action derived from the interaction between its current phenotype and the micro-environment conditions. For that, in a first decision, the intrinsic phenotype given by the random variable 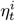has to be compared with a threshold value, 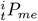. This threshold values determines if a cell retains or switches its phenotype. Since this is performed by the oxygen concentration and cell density, 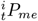 must be modeled as a function expressed in terms of Equations (4) and (6). Each function, 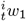or 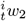, will act as a relative weight function on the other according to Equation (8) which is depicted in Figure 7. The denominator has been formulated in order to get *P*_*me*_ = 1 when either 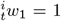or 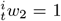. Since Equations (4) and (6) can only take values comprises between 0 and 1, 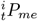 will also have values within this interval.

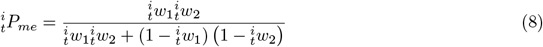

**Figure 7:**
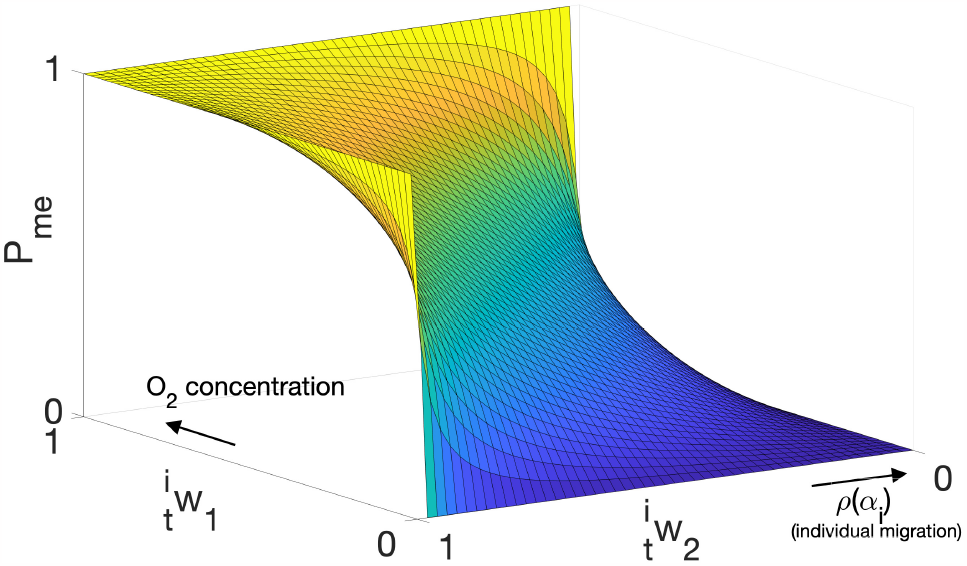
Graphical representation of the threshold function 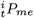, Equation (8). The arrows point the oxygen and cell density growth related to the values of 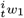 and 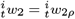, respectively.

For example, consider 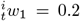and 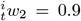. These values mean that the oxygen field in the micro-environment promotes the cell migration (low level of oxygen), 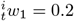, whereas the cell density promotes its quiescence, 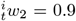. In order to make a decision, both values are weighted by Equation (8) resulting in *P*_*me*_ = 0.6923. The cell decision is the result of comparing the value of *P*_*me*_ to the value of intrinsic phenotype, given by 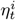. The decision process will be explained in Section 3.2.6.

#### 3.2.4 Boundary conditions

The boundary conditions of the ABM are a critical point in the formulation. In this model, instead of using a rigid boundary condition that corrects or prevents a certain movement, a soft boundary condition is implemented in the decision model through the information layer. At the information layer, the density field is forced to be *ρ* (*α*_*i*_) = 1 in the regions where a boundary condition is applied to restrict the movement. Indirectly, when the ABM model looks for the density in a region with a boundary condition the value of *ρ* (*α*_*i*_) = 1 it results *P*_*me*_ = 0, and so the cell will not tend to move towards a region with this boundary condition.

#### 3.2.5 Interaction between the cell and its adjacent micro-environments

The micro-environment conditions of adjacent elements of destination will condition the cell migration since the final action will depend on the oxygen and cell density conditions in the closer elements. By the assessing of the micro-environment in the surrounding elements, a preferential direction of migration can be obtained by Equation (9).

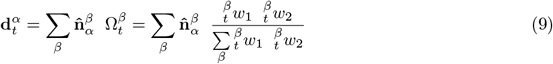

In this equation, the final migration direction 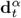, between the current element *α* and all possible destinations *β*, is calculated as follows. Each unitary vector 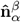, that connects every pair of origin-destinations elements, is weighted with a coefficient 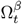. This coefficient represents the relevance that the micro-environment conditions of a certain destination *β* has with respect to the oxygen and cell density conditions of all elements of destination as a whole. It is calculated dividing the information relative to the oxygen concentration and the cells density in a certain element of destination, given by 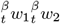, by the sum of this information in all possible element of destination. Once each pair of unitary vectors has been weighed by their corresponding coefficient 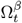, they are summed.

The Figure 8 shows a 2D central element *α* surrounded by eight possible elements of destination *β*. In this case *β* = 1, …, 8. The eight unitary vectors 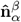 are depicted in green and dotted arrows. One of them has been drawn in solid arrow to highlight one of the possible unit direction of destination. The final direction of destination 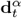 is represented in red.

**Figure 8:**
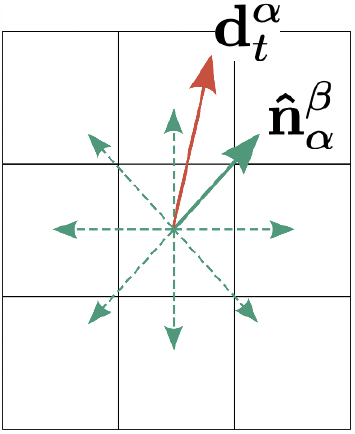
Scheme of the surroundings of a central element in 2D, *α*, with the eight possible directions of destination 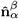, in green. The final one, 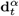, is depicted in red

The final displacement of the cell, 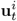, is obtained through Equation (10) multiplying the unitary vector of Equation (9), 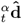, by the average velocity of the cell during its displacement, *v*_*i*_, and by the time increment, ∆*t*, in every time step, turning out

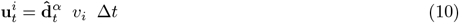

#### 3.2.6 Phenotypic plasticity modeling

Since the final phenotype adopted by a cell is directly related to its decision, the phenotypic plasticity will be modeled through a hierarchic decision structure. The process of decision, to stay or to migrate, of a single cell at each time step, ∆*t*, is accumulated up to a certain time *t* and includes primary decisions and subsequent ones derived from them. The cell primary decision is performed when its micro-environment conditions, given by Equation (8), allow that a cell to keep or to switch a intrinsic phenotype. This decision will also depend on the micro-environment of adjacent elements so that Equation (9) also plays an important role. The rest of phenotypes will be a consequence of secondary decisions taken by cells from the primary ones. To account all of these possibilities, the following hierarchic decision process has been designed adopting a structure of tree, Figure 9.

**Figure 9:**
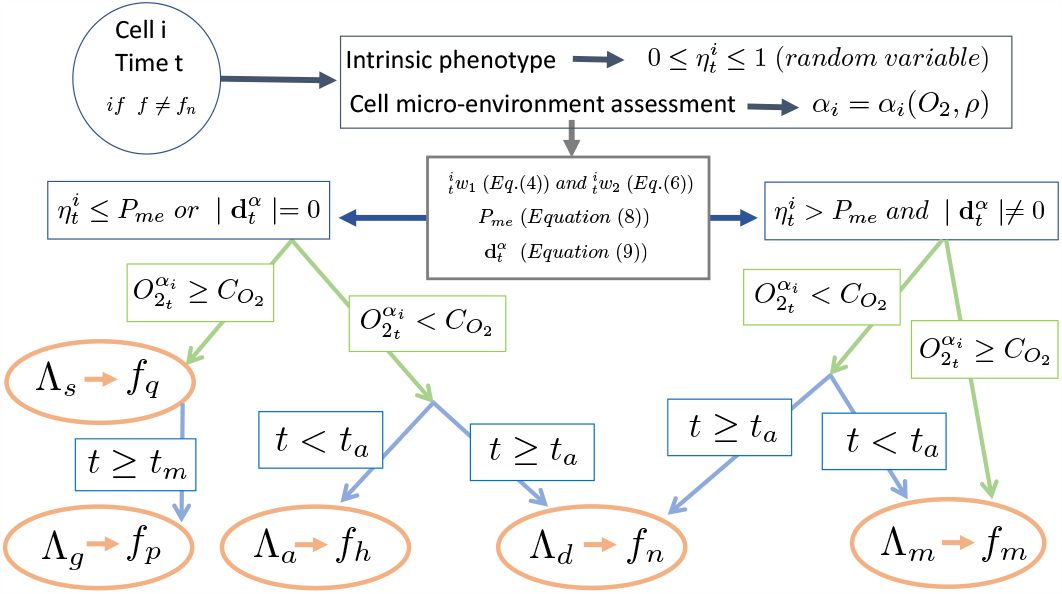
Hierarchic decision tree to model phenotype plasticity.

The first step is to check if the cell is alive or dead, blue circle in Figure 9. If it is dead, that cell will keep the necrotic phenotype and quit the decision process occupying its current position in the mesh. If the cell is alive, the intrinsic migratory or quiescent phenotype is determined by the random values of 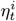 assigned to every cell at each time step. Since a low standard deviation will be adopted in 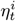, a little variation in the intrinsic phenotype will be considered at each time step.

##### Primary decisions

As above explained, if 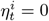, a cell will always tend to stay. In this case 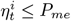 is always fulfilled, ∀*P*_*me*_. On the other hand, if 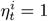, the cell will try to migrate. In this case 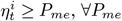. Therefore, for the rest of the values of 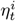, the primary cell decision will be to stay whenever 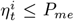 is fulfilled whereas the action of migrating will be executed if 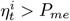. There is a case in which the cell may decide to stay when 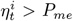. This is produced when there is no space in the neighbor elements and/or the oxygen available in destination is not sufficient. This is mathematically modeled through Equation (9) when 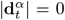 (a tolerance may be established here). Thereby, the primary decision of the cell will be to stay if 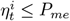 or 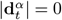 whereas it will be to migrate if 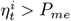 and 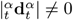 0, as it is depicted in Figure 9 in navy blue color. In the former, cell will adopt quiescent phenotype whereas in the latter a migratory one.

Then, some subsequent decisions are performed. In this point, genotypic variables are paramount in the decision of the cells.

##### Subsequent decisions for quiescent cells

When the primary decision is to stay, i.e. when 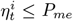 or 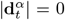, there are two alternatives for the cell, depicted in green in Figure 9. If the oxygen concentration in its micro-environment, 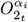, is greater or equal to the minimum oxygen level required by a cell, 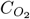, the final decision will be to stay. In this case, the final cell phenotype will be quiescent, *f*_*q*_, although it can be switch to proliferative, *f*_*p*_, if the time in quiescence status is greater or equal than the minimum required to produce mitosis, i.e. *t* ≥ *t*_*m*_. When 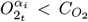, i.e. under anoxia conditions, there are two additional options which are depicted in blue in Figure 9. On one hand, if the accumulated time *t* under anoxia is greater or equal than the maximum time that a single cell can be in apnea *t*_*a*_, the final decision will be to die Λ_*d*_ and the phenotype will also be necrotic, *f*_*n*_. However, if *t < t*_*a*_, the final action will be to stay in apnea, Λ_*a*_ to adopt a hypoxic phenotype, *f*_*h*_.

##### Subsequent decisions for migratory cells

When the primary decision is to migrate, i.e. 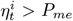 and 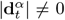, the secondary cell decision, in green in Figure 9, will also depend on whether the oxygen available in the element of destination is lower than the minimum concentration of oxygen needed by a cell to survive or not. If there exists enough oxygen the secondary decision of the cell will be to migrate. If the oxygen concentration is lower than the requested by a cell to survive two additional decision will be performed, in blue in Figure 9. On one hand, the cell will die, adopting a necrotic phenotype, if the time under hypoxic conditions is greater or equal than the maximum time that a single cell can be in apnea *t*_*a*_. On the other hand, if this time is lower, the cell will migrate in apnea, with a combined hypoxic/migratory phenotype.

#### 3.2.7 Simplification for grouping and for 2D cases

The above-explained model can be applied to both individual cells or clusters of cells. In the latter, all cells of the cluster will have the same genotypic properties and will decide as if they were a single cell. On the other hand, although the model was initially developed for performing 3D simulations, 2D cases can be executed considering the diameter of the cells or the cluster’s as the third dimension

## 4 Parametric study and sensitivity analysis

### 4.1 Influence of the mathematical parameters of the model

In this section, an analysis of the influence of the different values of *A*_1_, *B*_1_, *ρ*_*op*_ and *w*_*f*0_ in the cell behavior is addressed. Table 3 shows the figures in which the results of Equations (4), (5), (7), (6) and (8) are respectively depicted. The results of each equation are represented on the vertical axis. The equations are computed for different values of the variable at hand 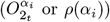, depicted on the left axis, and for different values of the parameters, on the right axis.

**Table 3:**
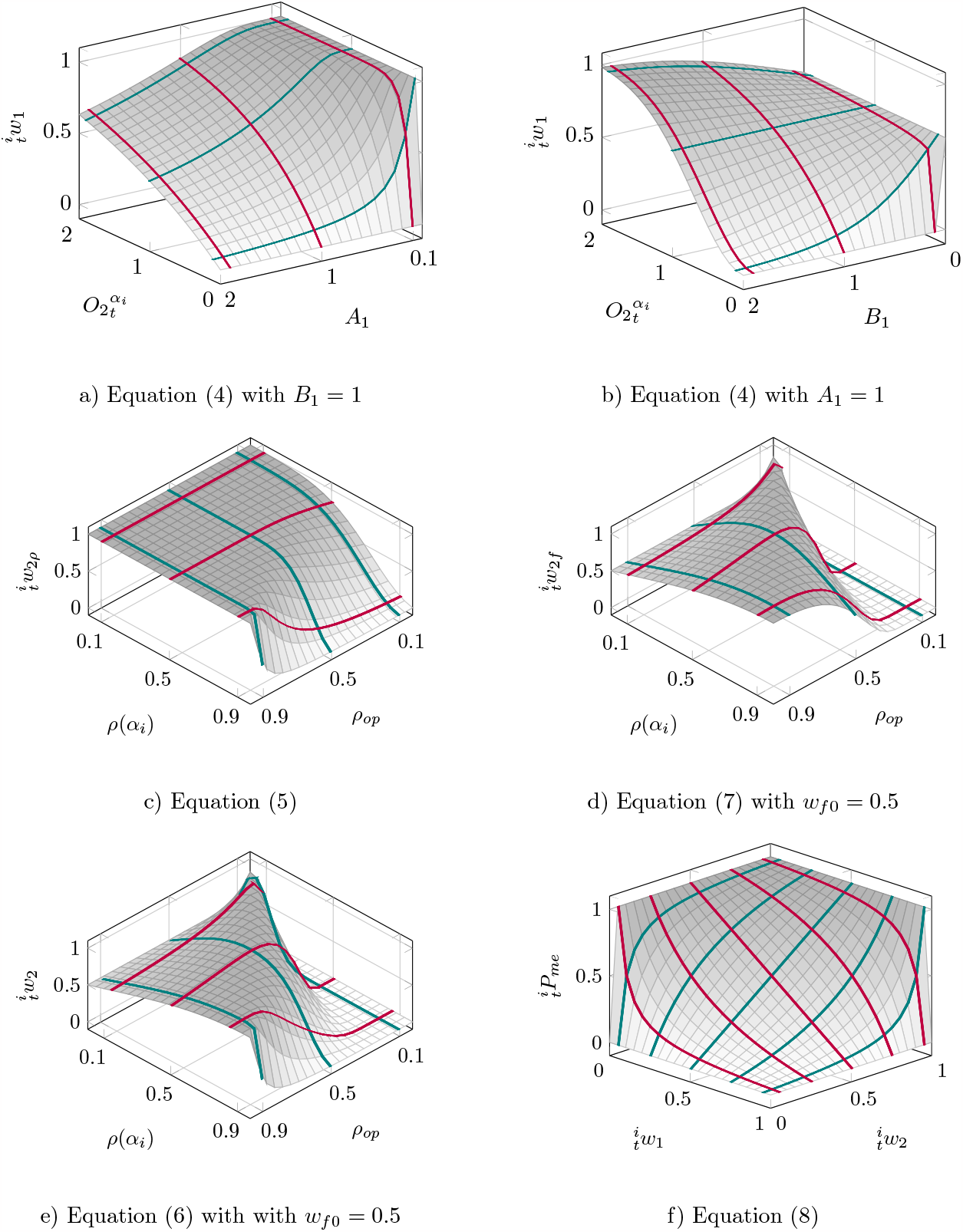
Graphic representation of the interaction between parameters in the equations used in the model. The green and red lines above the surfaces shows 2D cases of the equations presented, where the extreme and average values of one variable are fixed.

The results of 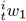, Equation (4), are shown in Figures a) and b). These figures show the influence of the parameter *A*_1_ (keeping *B*_1_ = 1 constant) and *B*_1_ (taking *A* = 1) in the behavior of a cell with a given oxygen level in its surroundings 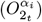. According to Figure a), for any oxygen concentration, the probability of a cell staying is increased as the values of *A*_1_ are lower (green curves). The same behavior is observed for *B*_1_ when the oxygen concentration is low, Figure b). However, for higher levels of oxygen, the probability of a cell staying is increased as *B*_1_ is higher. This change in the way in which *B*_1_ influences the cell behavior is produced when 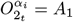, Figure b). Figures a) and b) also show that the influence of both parameters is greater for low oxygen concentrations.

The parameters that govern the collective behavior and the influence of the cell density in the behavior of a cell are, respectively, *w*_*f*0_ and *ρ*_*op*_. The influence of these parameters in the cell behavior is depicted in Figures c), d), and e). In these figures, the values of Equations (5), (7), (6) for different values of *ρ*_*op*_ and for *w*_*f*0_ = 0.5 are depicted. *w*_*f*0_ forces the probability of the cell staying is constant an equal to the value adopted for *w*_*f*0_ for any *ρ*_*op*_ when *ρ*(*α*_*i*_) = 0. Since we are always going to obtain that the probability is equal to *w*_*f*0_ when *ρ*(*α*_*i*_) = 0, we have adopted the only value of *w*_*f*0_ = 0.5 for the sake of clarity. In Figure c), it is observed that, in general terms, when *ρ*_*op*_ is decreased the probability of a cell staying is also reduced, being more evident for high values of the cell density, *ρ*(*α*_*i*_), red curves in Figure c). The influence of *ρ*_*op*_ in the probability of a cell staying is reduced when the cell density is lower. This makes sense, since low cell densities do not trigger cell migration. This behavior is corrected by the function 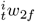whose results are depicted in Figure d). In this figure, it is observed that there is a maximum at *ρ*_*op*_ which means that the probability of a cell to stay is maximum when *ρ*_*op*_ is reached, so the cell will tend to migrate up to form cell clusters with a cell density equal to *ρ*_*op*_. In Figures d) and e), the graphical representation of the maximum values of both 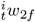 and 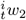 in *ρ*_*op*_ = *ρ*(*α*_*i*_) = 0 impedes from seeing the fact that 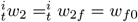 whenever *ρ*(*α*_*i*_) = 0.

Finally, Figure f) shows the results of Equation (8) in terms of 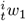 and 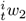. This figure is the same as Figure 8 and although 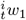 and 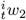 are not parameters, we consider important to outline that low values of 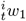 and 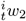 implies a nearly null probability of the cell staying and high values of them produces a high probability of the cell staying.

### 4.2 Sensitivity analysis of the main genotypic parameters

The influence, in the cell behavior, of the main genotypic parameters of Table 1, i.e. 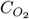, *t*_*a*_, *v*_*i*_ and 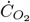, is addressed in this section. This study will be performed in the case of collective migration since it is predominant in GBM, see Section 1.

The analysis is performed by studying the evolution of an initial cell population consisting of 1000 cluster of cells distributed and aligned in a circumference of a radius of 1 mm, in a domain of 5.5×5.5 *mm*^2^ divided into 40×40 elements. The cells are considered spherical with a diameter of 0.025 mm. The oxygen in the domain at the beginning of the simulation is constant and with a value of 1 mmHg. The boundary conditions consist of a constant oxygen concentration of 7 mmHg in all the borders of the square and during all the computation, Figure 10.f. The evolution of this environment will be analyzed in different scenarios. A reference configuration is considered with the parameters shown in the second row of Table 4. Then, one of the parameters will be changed, adopting, respectively, the maximum and the minimum values given in Table 4, keeping the rest of them with the reference values. These maximum and minimum values do not mean limits or restrictions of the parameters. Apart from the reference parameters used, the cells have a time to proliferate of 5 days. A high trend of the cells to migrate collectively is considered in all cases adopting *w*_*f*0_ = 0.37. The total time simulated is of 10 days.

**Table 4:** Values of the main genotypic variables used in the analysis of Figure 10 and Figure 11.

| | $C_{O_2}$ [mmHg] | $t_a$ [days] | $v_i$ [mm/s] | $\dot{C}_{O_2}$ [mmHg/s] | $\rho_{op}$ [-] |
| --- | --- | --- | --- | --- | --- |
| Min | 1.0 | 0 | $10^{-10}$ | $10^{-14}$ | 0.2 |
| Ref | 1.8 | 1 | $10^{-6}$ | $10^{-9}$ | 0.7 |
| Max | 2.6 | 15 | $2 \times 10^{-6}$ | $10^{-5}$ | 1.0 |

**Figure 10:**
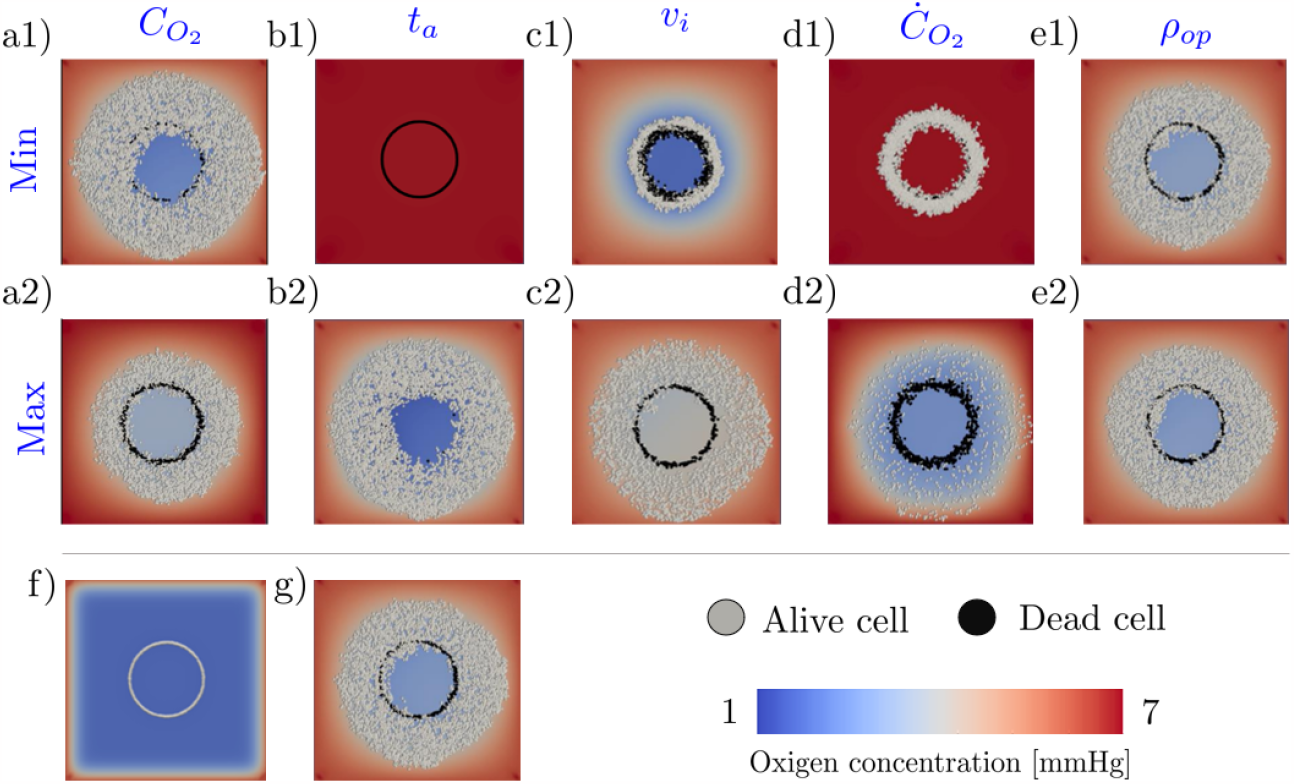
Evolution of the cell population and the oxygen field after 10 days. Rows 1 and 2 show the final configuration for the minimum and maximum values, respectively, of the genotypic variable indicated on the top of each column. f) and g) are the initial and final configurations of the reference case.

The results of the simulations are depicted in Figure 10. This figure shows the evolution of the initial cell population, Figure 10.f, after 10 days. The final configurations for the minimum and maximum values of the genotypic parameter at hand are respectively depicted in the first and the second rows. It can be appreciated that the results reproduce not only the proposed “go-or-stay” dichotomy but also the classical “go-or-grow” one.

The results of Figure 10 are quantified in Figure 11 in terms of alive and dead cells. Alive cells include the migratory, hypoxic, quiescent and proliferative phenotypes. In all cases, except for *t*_*a*_ = 0, the cells population is increased with respect the initial one of 1000 cells, because of the constant supply of oxygen from the boundaries. The exception is when the cell dies without a previous time in hypoxia, i.e. when *t*_*a*_ = 0. In this case all cells die at the beginning. This is produced because, for *t*_*a*_ = 0, the minimum oxygen concentration required by the cell to be in normoxia is 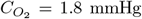. The initial oxygen available is 1 mmHg (*<* 1.8) in all the domain except in the borders. In this situation, cells would switch their phenotype to migratory but there is not enough oxygen available in the surroundings either. Then the phenotype should finally change to hypoxic, but since *t*_*a*_ = 0 all cells will die simultaneously, Figure 10.b1.

**Figure 11:**
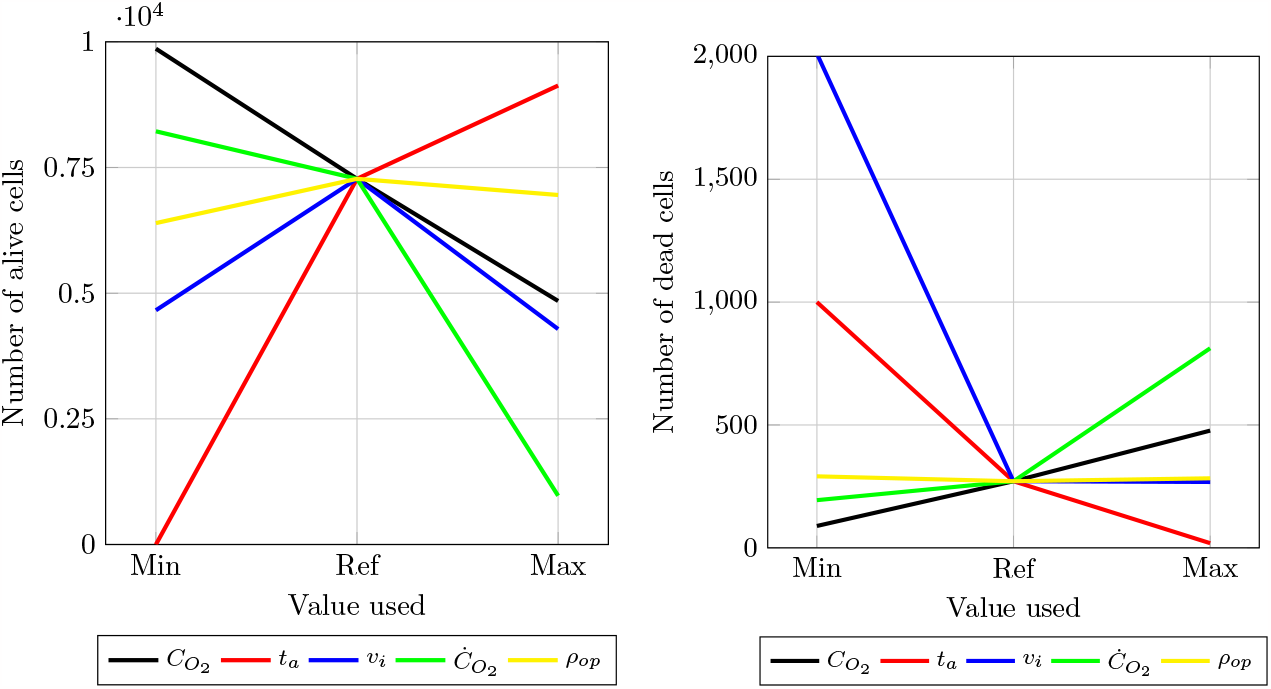
Results of the tests after 10 days, alive cells (left) and dead cells (right), for different values of the genotypic variables.

The reference value of the minimum oxygen concentration for not being in apnea was fixed in 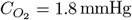 to force the cell migration. For the same reason, the minimum value is 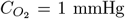, Table 4. According to the results depicted in black in Figure 11, the lower the values of the 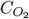 are, the more proliferation of cells is produced and the more cells survive. That means that if 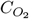 is low, cells will not migrate and after a time *t*_*m*_, they will proliferate. The new cells with this genotype will take more advantages than cells with a greater 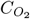 for the same oxygen concentration so they will survive longer. Consistently, the greater 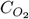 is, the more cells die and the lower cells survive, Figures 10.a1, 10.a2 and 11.

The results of the influence of the rate of oxygen consumption, 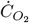, are depicted in green in Figure 11. According to these results, a lower rate of oxygen consumption implies more oxygen available in a certain period of time, promoting better environmental conditions for the survival of the cells, Figure 10.d1. The opposite thing happens when 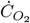is high, the oxygen will be depleted faster, promoting the cellular death, Figure 10.d2.

These results show how the high survival of GBM cells in hypoxic environments can be easily simulated through 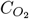 and 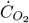.

Relative to the maximum time that a cell can be in apnea, *t*_*a*_, red lines in Figure 11, the lower *t*_*a*_ is, the more cells die, whereas the higher *t*_*a*_ is, the more cells live, Figures 10.b1 and 10.b2. As above explained, when *t*_*a*_ = 0 all cells die simultaneously.

The results corresponding to the average velocity of a cell during displacements, *v*_*i*_, are depicted in blue in Figure 11 (remind that we are considering collective migration). These results are very interesting. The minimum value considered for this parameter is *v*_*i*_ = 10^−10^ mm/s which means that cells migrate very slowly. This low velocity during migration produces a depletion of the oxygen in the area, Figure 10.c1, and consequently an increase of the dead cells compared to the results of the reference velocity, *v*_*i*_ = 10^−6^ mm/s. The results for the maximum velocity, *v*_*i*_ = 2 *·* 10^−6^, show fewer alive cells than in the reference case, keeping constant the number of dead cells, Figure 11. Therefore, the drop in the living cells is not because more cells die but because of a decrease in the cellular proliferation. This means that when a group of cells migrate with a suitable velocity, they can reach far areas with a higher oxygen concentration more easily, following the oxygen gradients, at the expense of not proliferating.

This prevalence of the survival of the cells that compose the group will be also found, in the following section, in the conformation of the necrotic cores in GBM cells with collective behavior.

This behavior is also observed in the results corresponding to the optimum density. In Figures 10.e1, 10.g and 10.e2, three similar final configurations are observed for the minimum, reference and maximum values adopted for *ρ*_*op*_, namely: 0.2, 0.7 and 1, respectively. This similarity is quantitatively justified in Figure 11 with the results depicted in yellow. According to these results, in the case of collective migration, no matter the size of the clusters, which is directly related to the values of *ρ*_*op*_, cells will migrate in groups prevailing the group survival over the proliferation which is aligned and with the previously commented for the case of velocity *v*_*i*_. The conclusions for *ρ*_*op*_ and *v*_*i*_ also fulfills both the “go-or-stay” and the “go-or-grow” dichotomies.

## 5 Cases of study and results

Aside the cases presented in the sensivity study, four more examples of application of the proposed model are shown. Each case reproduces different cell behaviors. This cell characterization has been modeled through the phenotype and genotype parameters shown in Table 5 which have been considered constant for all cells in each case. The intrinsic phenotype of the cells corresponding to the three first cases is mainly migratory, given by 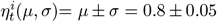, whereas for the fourth case there is not a predominant migratory or quiescent intrinsic phenotype since 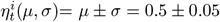.

**Table 5:** Phenotypic and genotypic parameters.

| Parameter | Units | C-1 | C-2 | C-3 | C-4 |
| --- | --- | --- | --- | --- | --- |
| $\mu$ | $[-]$ | 0.8 | 0.8 | 0.8 | 0.5 |
| $\sigma$ | $[-]$ | 0.05 | 0.05 | 0.05 | 0.1 |
| $B_1$ | $[-]$ | 0.01 | 0.01 | 0.01 | 0.05 |
| $A_1$ | $[mmHg]$ | 0.2 | 0.2 | 0.2 | 0.7 |
| $\rho_{op}$ | $[-]$ | 0.0 | 0.7 | 0.0 | 0.0 |
| $w_{f0}$ | $[-]$ | 0.99 | 0.37 | 0.99 | 0.99 |
| $\dot{C}_{O_2}$ | $[nmol/(cell \cdot min)]$ | $8.46 \cdot 10^{-4}$ | $8.46 \cdot 10^{-4}$ | $8.46 \cdot 10^{-4}$ | $3.06 \cdot 10^{-4}$ |
| $t_a$ | $[h]$ | ∞ | 20 | 20 | ∞ |
| $v_i$ | $[mm/s]$ | $1.38 \cdot 10^{-4}$ | $1.38 \cdot 10^{-4}$ | $1.38 \cdot 10^{-4}$ | $1 \cdot 10^{-6}$ |
| $C_{O_2}$ | $[mmHg]$ | 0.5 | 0.5 | 0.5 | 1.8 |
| $t_m$ | $[h]$ | ∞ | 0.125 | 0.125 | ∞ |

The parameters corresponding to C-1 represent the cell migration induced by oxygen chemotaxis and cell density. A high influence of the cell density in the cellular migration and no flocking behavior have been considered adopting *ρ*_*op*_ = 0 and *w*_*f*0_ ≈ 1, respectively. The option of necrotic and proliferative phenotypes have not been regarded by taking *t*_*a*_→ ∞ and *t*_*m*_→ ∞. In case C-2, a low sensitivity to high cell density, *ρ*_*op*_ = 0.7, and flocking behavior, *w*_*f*0_ = 0.37, are included together with the oxygen chemotaxis. In order to study the behavior of these cells but without the flocking effect, C-3 case has been regarded. The formation of necrotic cores with C-2 and C-3 cells behavior has been also addressed. In the fourth model, C-4, the parameters have been estimated to reproduce a set of experimental results. Specifically, the C-4 case simulates the cells migration induced only by cell density gradients, in a micro-channel.

The values of the genotypic parameters can be obtained from experimental tests available in the literature. However, the results of these experimental measurements show a heterogeneity and fluctuation depending on the conditions and tumor regions. Such is the case of the oxygen concentration in GBM, where intertumoral areas are less oxygenated than peritumoral ones [13, 4, 18]. This heterogeneity is due to the different oxygen consumption rates since glioma cells in oxygenated zones consume more oxygen than glioma cells in hypoxic zones, [4]. Relative to the oxygen consumption rates there is also heterogeneity in the results of experiments. For example, the oxygen consumption rate in SF188 glioma cells varies, depending on the substrate, between 0.15 and 0.25 *nmol/min* [53] whereas in U251 astrocitoma cells, the oxygen consumption rate is given between ≈ 2.4 *·* 10^−6^ and ≈ 4 *·* 10^−6^ *nmol/min* [33]. Relative to the hypoxic conditions induced by the oxygen consumption, there is a homogeneity to denominate “poorly oxygenated” areas when the oxygen concentration is less than 5 *mmHg* which means less that 0.7% of *O*_2_ in the gas phase. An area is called “modestly hypoxic” when oxygen concentrations is given between 0.6% and 2% of *O*_2_ [4, 18, 62]. Hypoxic conditions also alter the proliferation parameters which can be estimated from cell population doubling times. McCord et al. found that mean population doubling times for GBM-stem-cells in neurosphere cultures under 20% of *O*_2_ turned out between 118 and 35 hours, whereas they were significantly reduced to 80.36 and 26.29 hours for cultures under 7% of *O*_2_ [48].

All these considerations are related to the genotypic parameters of the present model. Thus, we have adopted 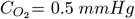 for C-1, C-2 and C-3 which is in agreement with the values adopted by [47] since in our cases there is no oxygen supply during all the simulation. In C-4 the level of oxygen is kept constant and therefore 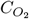 is not relevant whenever 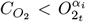. The oxygen consumption rate, 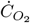, has been adopted between the values given by Griger et al. [33] and Pike et al. [53] in all cases. Relative to the time needed to produce mitosis, single cell doubling time is *t*_*m*_= 0.125 hours per cell in those cases where proliferation is produced. The value of the averaged velocity of the cell during displacement are based on the results of Yan and Irimia [65]. Finally, the maximum time in hypoxia before cell death, *t*_*a*_, was selected for convenience. Details of the influence of this parameter can be seen in Section 4.

All cases have been applied to bi-dimensional domains according to the specified in Section 3.2. In all of them, boundary conditions of *ρ* = 1 has been applied on the boundaries of each domain in the discrete layer whereas 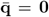 was imposed on all boundaries in the continuum one. In all cases, both individual cell and cell clusters are considered spherical. After running these examples and the cases of the sensitivity study, Section 4, in a computer in a serial code of a core i5 CPU, the average runtime was 0.2 seconds per cell and time step.

### Case 1. Individual cell migration induced by oxygen and cells density

This example shows a population of 200 individual cells randomly disseminated in a domain of 13x13 *mm*^2^ meshed with a element size of 1 *mm*. Cells are characterized by the phenotype and genotype given by the parameters corresponding to C-1 in Table 5. The diameter of each cell is 0.022 mm.

Two cases of oxygen distribution, with no external income during the process, have been adopted. In the first case, the oxygen is initially modeled with a Gaussian distribution centered in the middle of the field, with a maximum of 0.7 *mmHg* and a radius of 3 *mm*. In the second oxygen field, the same distribution is used but centered in the right lateral and with a radius of 6 *mm*.

The results of the simulation are shown in Figures 12 and 13, respectively. The maximum concentration of oxygen is represented in red whereas the minimum is depicted in blue. In order to clearly understand the migration process and the evolution of the oxygen field, only three phenotypes have been considered, namely quiescent, migratory and potentially proliferative. Although mitosis has not been regarded in these cases, the amount of cells depicted in white could increase because of the time in quiescence and the oxygen and the cell density conditions. This phenotype can be renamed as potentially proliferative in this case. The cells with quiescent phenotype are shown in yellow whereas the ones with a migratory phenotype are depicted in green.

**Figure 12:**
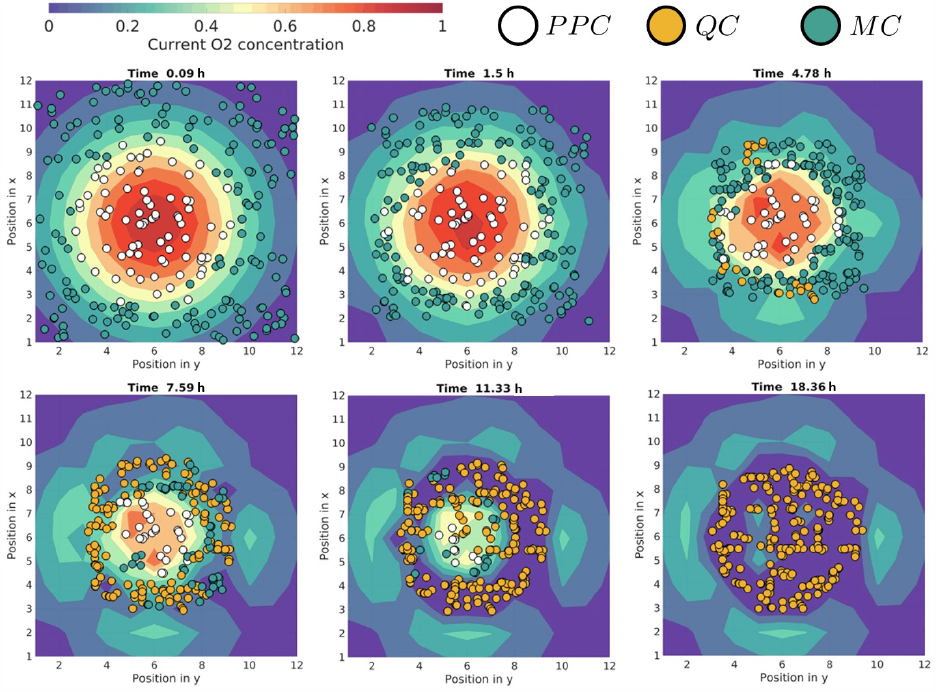
Individual migration. Evolution of the oxygen field and phenotypic plasticity during migration. Centered gaussian distribution of oxygen. PP: potentially proliferative cells, QC: quiescent cells, MC: migratory cells.

**Figure 13:**
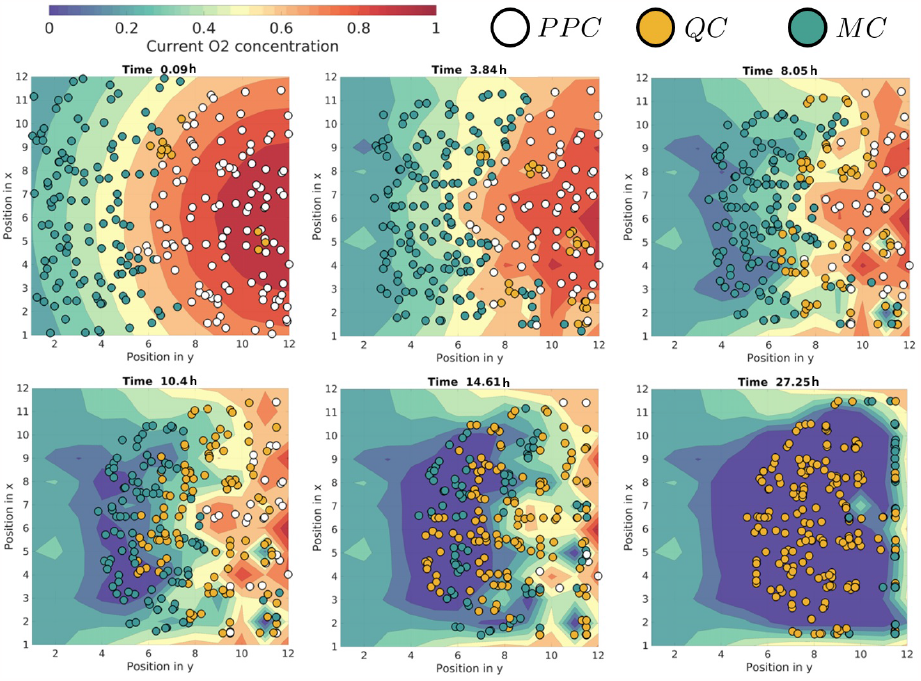
Individual migration. Evolution of the oxygen field and phenotypic plasticity during migration. Lateral gaussian distribution of oxygen. PP: potentially proliferative cells, QC: quiescent cells, MC: migratory cells.

In both figures, it is observed a migratory process from zones with the lowest oxygen concentration towards areas with the highest level of oxygen, as in the experimental microfluidic tests [8]. The field of oxygen is simultaneously evolving so that, in the end, most of the oxygen available has been consumed by the cells. Although in the two simulations the figure shows that the initial intrinsic phenotype is markedly migratory, a switch to quiescent and/or to potentially proliferative ones is produced depending on the oxygen available and the accumulation of cells in certain zones. In Figure 12, all the cells have a migratory or potentially proliferative phenotype up to the third frame. Potentially proliferative cells are concentrated in zones with the highest oxygen concentration. Migratory cells are placed in areas with the lowest oxygen concentration. From the third frame, the phenotypes of the cells start to evolve to a quiescent one due to the restriction of the cells density and the low oxygen concentration in their surroundings. At the end of the process, all cells adopt this quiescent phenotype and are concentrated around the zone where there is more oxygen available. At this moment, the field of oxygen is almost constant and close to zero which makes cells be potentially necrotic. A similar behavior is observed for the case depicted in Figure 13.

### Case 2. Collective migration induced by oxygen and cell density signals

In Figure 14, we show the collective behavior of an initial population of 90 clusters composed, each one, by four cells (the diameter of each cell is 0.022 mm) in a domain of 3×3 *mm*^2^ which has been meshed with a square element size of 0.15 *mm*. The cells genotype correspond to C-2 in Table 5. To simulate the collective behavior, flocking between clusters has been included through the parameters *ρ*_*op*_ = 0.7 y *w*_*f*0_ = 0.37. As above explained, each cluster behaves as an individual cell. Only three groups of cells corresponding to three phenotypes have been depicted for sake of clarity. Migratory cells are shown in green, proliferative ones in orange and necrotic cells in black.

**Figure 14:**
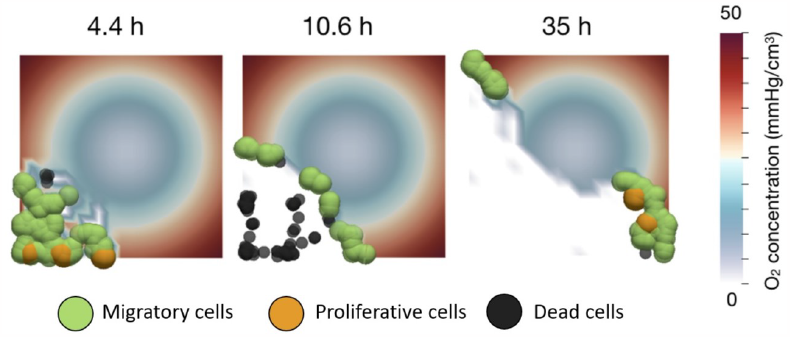
Collective migration. Evolution of the oxygen field and phenotypic plasticity during migration.

The initial population is positioned in the bottom left corner. The initial oxygen distribution is an inverse exponential centered in the middle of the plane. This distribution makes the greatest concentration of oxygen, depicted in red in Figure 14, be distributed along the sides of the plane whereas there is a lack of resources just in the center, depicted in blue color. When the cells deplete the oxygen available in the initial zone, they migrate forming more clusters of cells and following the paths with higher levels of oxygen. Therefore, the cells avoid the zone with the lowest oxygen concentration. The figure shows how the clustres die due to a prolonged apnea when they have no access to oxygen and cannot migrate. These cells and the new ones resulting from mitosis have been eliminated in the figure for sake of clarity.

### Case 2 and case 3. The necrotic core. Influence of the collective migration in its evolution

The evolution of the necrotic cores corresponding to two different cells population is shown in Figure 15. The only difference between them is that a flocking behavior has been considered for the case depicted at the bottom. For that, the parameters corresponding to C-2 in Table 5 have been adopted whereas the corresponding ones to C-3 have been considered for the cell population depicted at the top. In both cases, dead and proliferative phenotypes are also possible through *t*_*a*_ = 20 hours and *t*_*m*_ = 0.125 hours. Cells with proliferative phenotype are depicted in orange, whereas the migratory and dead phenotypes are depicted in green and black, respectively. The hypoxic and quiescent phenotypes has not been depicted for sake of clarity.

**Figure 15:**
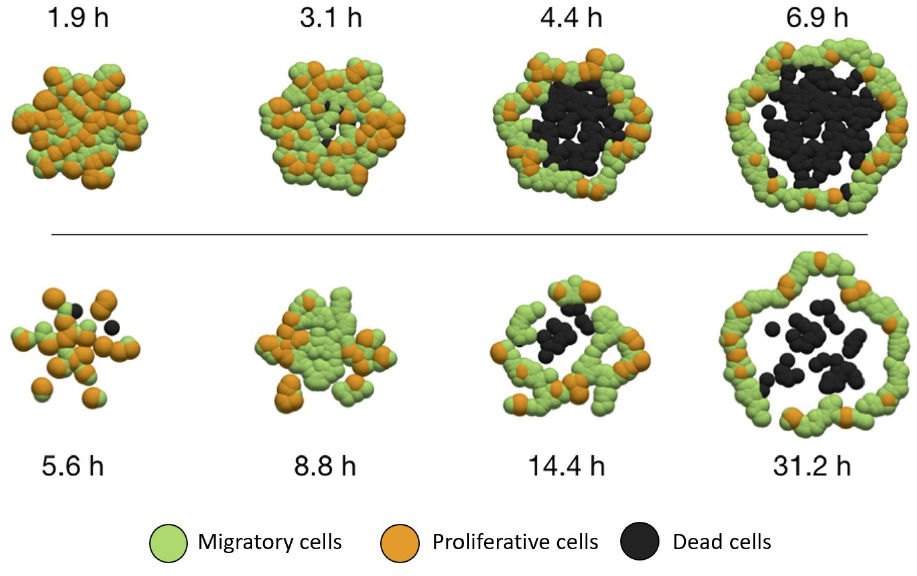
Necrotic core formation. Top) tumor cells evolution without flocking. Bottom) tumor cells evolution with flocking.

The initial cell distribution consists in 100 cells concentric and radially distributed from the center of a domain of 3×3 *mm*^2^. This domain was meshed with a square element size of 0.15 *mm*. In Figure 15, only the necrotic core surroundings have been depicted. There is no external income of oxygen in the field and the test starts with an uniform oxygen distribution. Since the results of the evolution of the oxygen field show the same trend as the ones previously described for C-1 and C-2, this has not been either included in the figure for sake of simplicity.

The necrotic core is formed after the cells consume the oxygen available in their locations. At this moment, the phenotype of the inner cells change to hypoxic since they can not migrate because of they are surrounded by many cells. Therefore, these inner cells are left without any oxygen supply so that they die after some time in apnoea. The outer cells form a ring of proliferative and migratory cells. As it can be deduced from Figure 15, the model is capable of reproducing the formation of the necrotic core in the cases considered. However, the cells with flocking tendency develop a less dense necrotic core and a fewer number of new cancer cells. Furthermore, the time invested in forming the necrotic core is clearly greater in the case of collective migration. The opposite thing happens in individual migration, cells prefer to proliferate instead of grouping together. Since the number of cells is increased with the time in the later case, the demand and consume of the oxygen will be also greater and therefore faster, so that the necrotic core is produced before when compared to the case of collective migration.

These results are quantified in Figure 16 where the evolution of the amount of death, proliferative and migratory cells are shown in function of the time. The results for the population of cells with individual migratory behavior are depicted in continuous curves whereas the ones for the cells with collective behavior are shown in discontinuous streak. The curves depicted in blue, read and green represent the number of proliferative, migratory and death cells, respectively. The total amount of cells are obtained adding the corresponding values of all these phenotypes and are represented in black color. In the first stages, there only exists proliferative cells when they do not flock whereas all kind of phenotypes are found in populations whose cells tend to flock. In the first case, there is a clear proliferation of cells until they start to migrate before one hour. In this moment the number of new cells descends. Before 3 hours, the oxygen available is not enough and some cells start to die reducing the total amount of cells. In the case of the population with flocking behavior, the number of dead and proliferative cells is almost constant until the end of the process. However, the amount of migratory cells increases constantly.

**Figure 16:**
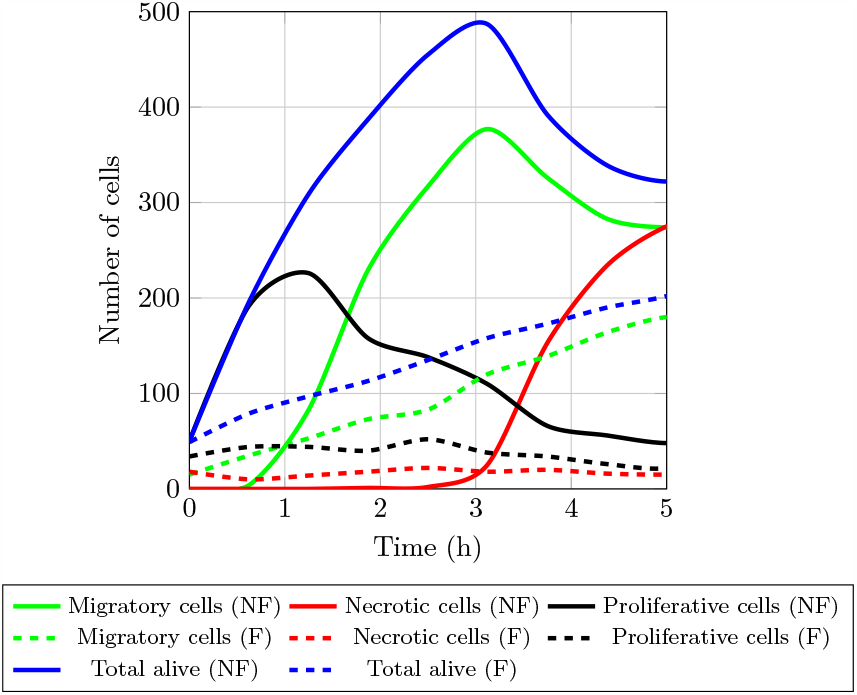
Evolution of the number of proliferative, migratory, dead and total cells during the necrotic core conformation. F: population cells with flocking behavior. NF: population cells with no flocking behavior.

This shows that cells with flocking behavior prioritize the creation of clusters of cells to the detriment of their proliferation. When there is a lack of oxygen in the environment, the cell clusters migrate increasing their amount up to reach the optimal density. This collective behavior increases the cells surveillance since it does not create an additional need of oxygen produced by cells proliferation. This is in agreement with previous studies where it was observed that pseudopalisading cells are between 5% and 50% less proliferative and 6 to 20 times more apoptotic than adjacent tumor areas, [54, 16].

Summarizing, cells with individual behavior migrate and proliferate before and faster than the cells with tendency to form clusters. However, the number of dead cells is lower in cells with flocking behavior since the collective survival is prioritized in this case.

### Case 4. Migration induced by cells density gradients

Marel et al. [46] studied the diffusion of cells in confined micro-channels considering the cell density as the only cue to produce the migration. They created a micro-channel on a petri dish so that its entry was continuously fed with cells. Authors used a particle image velocimetry to track the cells trajectories as well as to determine the velocity field and cells density. They observed that the time evolution of the cell density throughout the micro-channel was similar to a wave that, after a certain time, reaches a steady state characterized by a constant shape of the cell density curve.

In order to reproduce these results, the phenotypic and genotypic variables corresponding to the C-4 case in Table 5 were adopted. Additionally, a constant level of oxygen in the micro-channel was kept during all the simulation, so 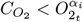. Thus, chemotaxis for oxygen gradients was not induced at the same time that cells had enough oxygen available. Keeping the oxygen concentration constant, ^*t*^*w*_1_ is also constant during the migration, Equation (4), so that Equation (8), and therefore the cell behavior, only depends on the cell density through the function 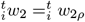, Equations (6).

The most difficult task for this simulation was to model the boundary conditions at the channel entry. We finally assumed that the cell density at the entry of the channel was constant. Thereby, when cells move, if the density at the entry is less than the initial one, cells are added in order to keep the density constant. For the initial setup, cells are distributed uniformly at the entry of the channel.

The model presented here does not include the cohesive forces between cells and, therefore, the simulation does not consider this effect. Despite this, the results shown in Figure 17 are promising. In this figure, the evolution of the cell density in three moments of the tests are shown. Experimental results have been depicted in dotted lines whereas the numerical predictions are represented by continuous lines. As it can be seen, the model reproduces the effect of the wave propagation shown in the experimental tests. Furthermore, there is also a good agreement between experimental results and model predictions which demonstrate the model possibilities to simulate migration triggered by density gradients.

**Figure 17:**
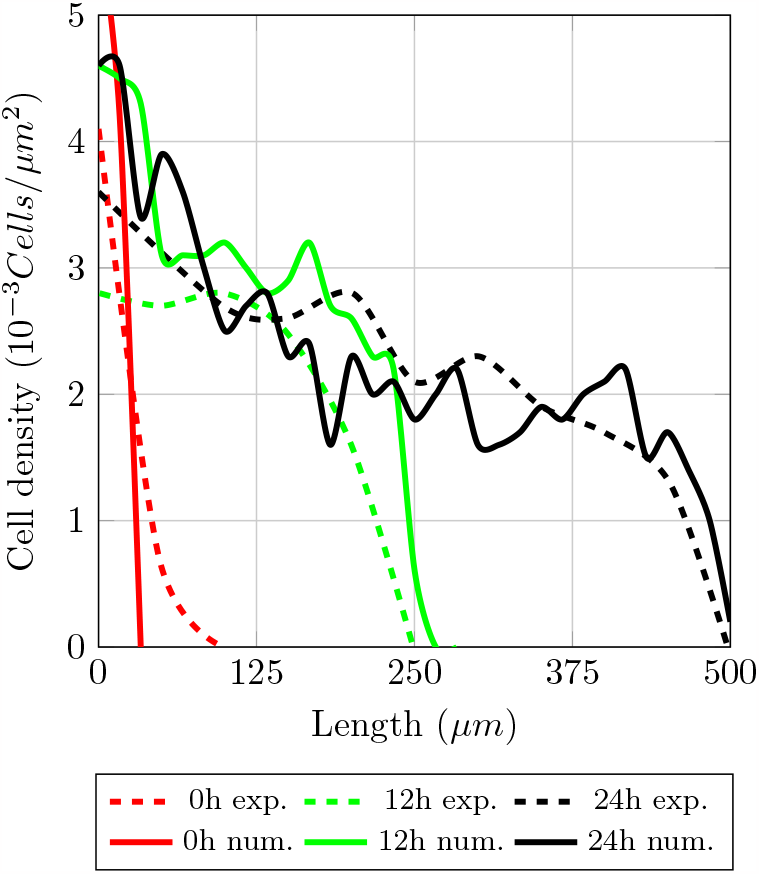
Experimental results and model predictions of cell migration in micro-channel.

## 6 Limitation of the model and future work

Despite the good results shown in the previous section, some limitations should be stated. The first one is the fact that sprouting angiogenesis has not been included in the model. Since the authors have developed a model to simulate of 3D angiogenic process in GBM [12], the next step would be to integrate both formulations in order to properly reproduce more realistic simulations. The second limitation concerns the fact of not including the forces between the cells. As in the case of the effects of angiogenesis, the effect of these forces can be indirectly considered through the parameters of the model, for example, *v*_*i*_. This would compel the calibration of these parameters with experimental tests which not always are available. The last limitation is that only isotropic migration is considered. This is related to the continuum layer that should be also improved with a more sophisticated formulation that includes, for instance, source terms that reproduce the oxygen supply during sprouting vasculogenesis.

## 7 Conclusions

A hybrid agent-based and continuum model has been presented to simulate the coupling between different multiscale events in GBM, such as the evolution of the oxygen field and phenotypic plasticity, during individual and collective cellular migration. The model is capable of reproducing five phenotypes, namely migratory, proliferative, quiescent, hypoxic and death, as well as the switch between them. The change between phenotypes is a consequence of the interaction between cells and their micro-environmental conditions. The behavior of the cells is characterized by genotypic parameters which have bio-physical meaning. The cell micro-environment is made up of the field of oxygen, which is simulated in the continuum part of the model, and the cell density, available in the ABM layer. Important bio-physics process have been successfully simulated, such as the cell migration, the flocking behavior of the cells and the conformation of necrotic cores.

A parametric and sensitivity analysis have been performed and four applications have been presented to test some of the model possibilities. The multiscale simulation has allowed to reproduce graphically and quantitatively the influence of both the oxygen concentration and the cell density in the phenotype of the cells. In all cases, the computation runtimes were good.

In the case of tumors whose cells present flocking, the model predicts that flocking behavior prevails over the cells proliferation regardless the size of the clusters. This implies that the oxygen is consumed at a slower rate than in the case of tumor cells without flocking since, in the later, new cells are produced and the oxygen demand is therefore increased. As a consequence of that, the necrotic cores of tumors with flocking cells are less dense and develop at a slower rate than in the case of tumor cells without flocking behavior.

Finally, the model has been applied to reproduce the experimental results of cells migration in confined micro-channels. In these tests, the only cue to migrate has been the cell density. According to the simulation results, the model is capable of reproducing the pattern of cells forming a wave throughout the channel that was found in the experimental results. Along time, the experimental results show an effect of a wave propagation which is also reproduced by the model.

The proposed model is promising for simulating many aspects of the GBM in order to increase our understanding of tumor evolution and answer “what if” questions in helping for such understanding.

## Acknowledgments

Grant PID2021-126051OB-C43 funded by MCIN/AEI/ 10.13039/501100011033 and by “ERDF A way of making Europe”.

---

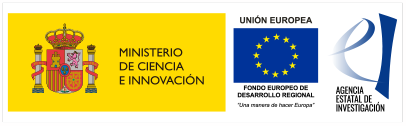

